# Sesbanimide R, a novel cytotoxic polyketide produced by magnetotactic bacteria

**DOI:** 10.1101/2020.12.22.424014

**Authors:** Ram Prasad Awal, Patrick A. Haack, Chantal D. Bader, Cornelius N. Riese, Dirk Schüler, Rolf Müller

**Affiliations:** Department of Microbiology, University of Bayreuth, Universitätsstraße 30, 95447 Bayreuth, Germany; Department Microbial Natural Products, Helmholtz-Institute for Pharmaceutical Research Saarland (HIPS)-Helmholtz Centre for Infection Research (HZI), German Center for Infection Research (DZIF, Partnersite Hannover-Braunschweig) and Department of Pharmacy, Saarland University Campus E8.1, 66123 Saarbrücken Germany

**Keywords:** Glutarimide-containing polyketides, cytotoxic activity, *trans*-AT polyketide synthase, magnetotactic bacteria

## Abstract

Genomic information from various magnetotactic bacteria suggested that besides their common ability to form magnetosomes they potentially also represent a source of bioactive natural products. By using targeted deletion and transcriptional activation, we connected a large biosynthetic gene cluster (BGC) of the *trans*-AT PKS type to the biosynthesis of a novel polyketide in the alphaproteobacterium *Magnetospirillum gryphiswaldense.* Structure elucidation by mass spectrometry and NMR revealed that this secondary metabolite resembles sesbanimides which were very recently reported from other taxa. However, sesbanimide R exhibits an additional arginine moiety the presence of which reconciles inconsistencies in the previously proposed sesbanimide biosynthesis pathway when comparing the chemical structure and the potential biochemistry encoded in the BGC. In contrast to sesbanimides D, E and F, we were able to assign the stereocenter of the arginine moiety experimentally and two of the remaining three stereocenters by predictive biosynthetic tools. Sesbanimide R displayed strong cytotoxic activity against several carcinoma cell lines.

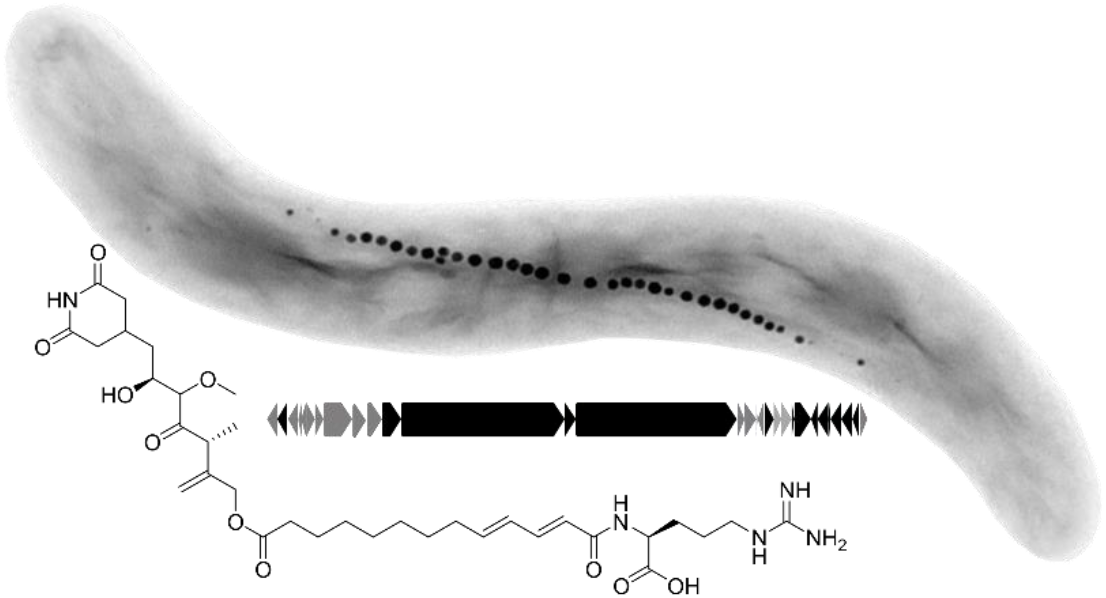

**Importance:** The finding of this study contributes a new secondary metabolite member to the glutarimide-containing polyketides. The determined structure of sesbanimide R correlates with its cytotoxic bioactivity characteristic for members of this family. Sesbanimide R represents the first natural product isolated from magnetotactic bacteria and identifies this highly diverse group as a so far untapped source for the future discovery of novel secondary metabolites.

## Introduction

Magnetotactic bacteria (MTB) share the ability to biomineralize membrane-enclosed organelles consisting of either magnetite (Fe3O4) or greigite (Fe3S4) crystals, called magnetosomes, which enable the cells to navigate within the Earth’s magnetic field (1). Studies on MTB so far have focused mainly on understanding magnetosome structure, biosynthesis and biological function as well as exploring the potential utility of magnetosomes as magnetic nanoparticles for various applications such as magnetic imaging or magnetic hyperthermia, magnetosome-based immunoassays, as nanocarriers in magnetic drug targeting and multifunctional nanomaterials with versatile functional moieties (2–7).

Apart from their common ability to form magnetosomes, MTB represent a highly heterogeneous group of prokaryotes. They are abundant and widespread in the sediments of many diverse aquatic ecosystems, ranging from freshwater to hypersaline habitats (8, 9), and besides a multitude of free-living, single-celled MTB, multicellular and even ectosymbiotic members of this group have been discovered (10–12). MTB are known to have diverse and versatile lifestyles, and members of this group are found in many different classes of eubacteria (13–15). Within the last years, a wealth of genomic information has been obtained by conventional, metagenomic, and single-cell genomics (16–20, 14, 15). We have recently shown that chances for the discovery of novel secondary metabolites clearly correlate with the increasing phylogenetic distance of the microorganisms under study (21). Because of their huge ecological, metabolic, phylogenetic and genomic diversity, producers of such interesting natural products might also be expected among MTB. Indeed, Araujo et al. (22) first noted the presence of typical secondary metabolite biosynthetic gene clusters (BGC), such as putative polyketide synthases (PKSs) and non-ribosomal peptide synthetases (NRPSs), in the genomes of several MTB. However, this so far has remained an untapped source for discoveries, largely owing to the fact that most of these bacteria are not tractable, or even cannot be grown at all in the laboratory.

One of the few MTB that can be cultivated reasonably well and is genetically tractable is the alpha-proteobacterium *Magnetospirillum gryphiswaldense* (23–26), which previously served as a model in many studies on magnetotaxis, organelle biosynthesis, and magnetite biomineralization (27, 2). Interestingly, also several putative BGCs for secondary metabolites were tentatively predicted in its genome (22, 26). This prompted us to investigate in more detail the strains’ biosynthetic capability using a combination of molecular and analytical methods.

In this study, we focus on the role of a *trans*-AT PKS BGC in *M. gryphiswaldense,* which we identify as a homologue of the sesbanimide gene cluster described by Kačar et al. in parallel to our studies (28). We set out to unambiguously assign the corresponding secondary metabolite from *M. gryphiswaldense* by markerless deletion of the gene cluster, to isolate the polyketide product and to elucidate its structure. Furthermore, we devise a model for sesbanimide biosynthesis that complements the one suggested by Kačar et al. (28) and reveals the new sesbanimide R as a missing link between the sesbanimide biosynthesis pathways, when compared across several taxa (29). In addition, we demonstrate cytotoxic activity of the novel sesbanimide congener.

## Results and Discussion

### Identification, deletion and transcriptional activation of a *trans*-AT PKS gene cluster

Using the antiSMASH tool (30), we identified several secondary metabolites gene clusters in the *M. gryphiswaldense* genome. Three gene clusters were predicted to encode the biosynthesis of a putative lasso peptide, an aryl polyene, and a homoserine lactone (Table S1 in the Supporting Information). In addition, a large (69,942 bp) gene cluster was predicted to encode a putative *trans*-AT PKS. It has a conspicuously high G+C content (66.7% vs 63.2% of the entire genome) and comprises 30 ORFs, which were tentatively assigned to various constituents of a *trans*-AT PKS.

To study the function of the cluster, we deleted the three putative core-biosynthetic genes (MSR-1_15630-15650) encoding two large polyketide synthases (PKS), a monooxygenase, and a gene (MSR-1_15620) encoding an acyltransferase. The deletion comprised 41,295 bp and yielded strain Δ*trans-at-pks* (Fig. S1 in the Supporting Information). Growth of strain Δ*trans-at-pks* was essentially wild type-like, with slightly increased doubling times during growth under aerobic conditions (Fig. 1A). Mutant cells were indistinguishable from the wild type with respect to cell length and shape (Fig. 1B, C, D, E). Cultures of Δ*trans-at-pks* exhibited a lower magnetic response (*C*_mag_=1.17, wild type: *C*_mag_ = 1.3; i.e. a light-scattering parameter for the semiquantitative estimation of average magnetic alignment of cells (31)). Transmission electron microscopy (TEM) of wild type (Fig. 1D) and Δ*trans-at-pks* (Fig. 1E) cells showed that both strains formed magnetosomes in about same numbers and with similar average sizes (Fig. 1F, G), however, both smaller (<25 nm) and larger (>60 nm) particles were more frequent in Δ*trans-at-pks* strain as compared to the wild type (Fig. 1F), which might explain the slightly lower magnetic response.

**Fig. 1:**
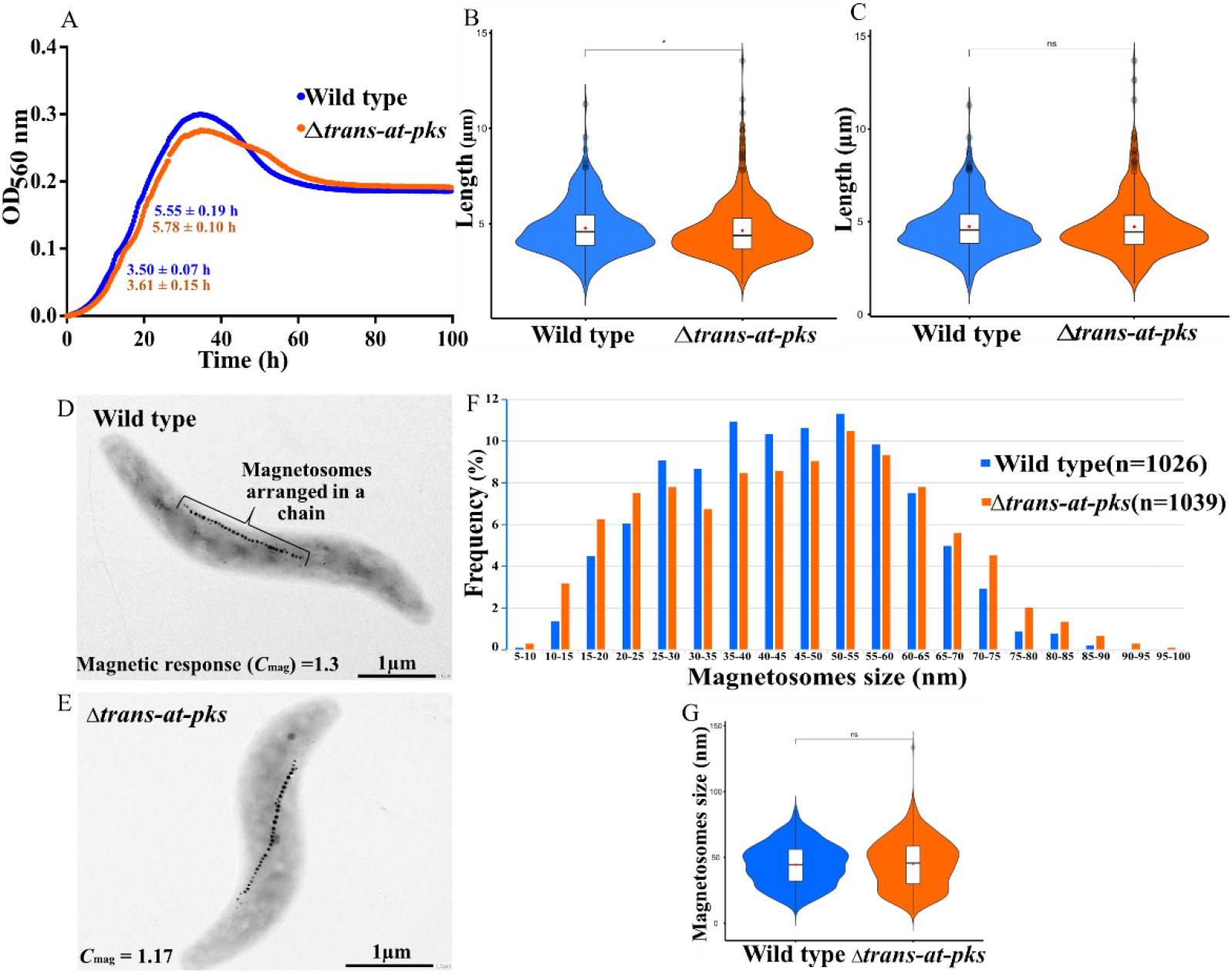
(A) Growth of the wild type and Δ*trans-at-pks* strains under aerobic conditions where the target compound was produced. Each growth curve represents the average of two individual growth curves. The doubling time (*Td*) (mean±SD) for each strain is given in the graph for the first and second part of the diauxic growth curve. (B) Cell length of wild type (mean=4.77±1.37 μm, n=312) and Δ*trans-at-pks* (mean=4.64±1.46 μm, n=504) grown under aerobic conditions. (C) Cell length of wild type (mean=4.73±1.37 μm, n=347) and Δ*trans-at-pks* (mean=4.72±1.6 μm, n=354) grown under microaerobic conditions. TEM images of wild type (D) and Δ*trans-at-pks* (E). Analysis of magnetosomes size distribution in wild type (mean=44.45±15.59 nm, n=1026) and Δ*trans-at-pks* (mean=45.18 ±18.29 nm, n=1039) (F, G).

To identify the biosynthetic product(s) of the *trans*-AT PKS cluster, wild type and Δ*trans-at-pks* strains were cultivated under aerobic, microaerobic and anaerobic conditions in FSM medium, and extracts of these strains were compared using principal component analysis (Fig. S2 in the supporting information) as previously described (32). Under microoxic and anoxic conditions, which are known to favour magnetosome biosynthesis (33, 34), there were no significant differences detectable between the mutant and the wildtype. However, in the extract of the wild type strain grown under aerobic conditions that are known to inhibit magnetosome formation (33, 34), we identified a compound with a mass of 691.38 Da which was absent from the mutant strain Δ*trans-at-pks.*

Yields of the target compound obtained from wild type cultures proved insufficient for the isolation and subsequent elucidation of its structure by NMR. Since we hypothesized that the low production might be due to poor expression of biosynthetic genes, we attempted to enhance their expression by transcriptional activation. To this end, a DNA fragment of 145 bp harboring a putative native promoter in front of MSR-1_15600 (ORF7) was replaced by a 64 bp fragment containing the stronger constitutive promoter *P_mamDC45_* (35) and the optimized ribosomal binding site (oRBS), yielding strain *P_mamDC45_-trans-at-pks* (Fig. S3 in the Supporting Information). P_*mamDC*45_ is an optimized version of the native promoter P_*mamDC*_ which drives transcription of the *mamGFDC* operon involved in magnetosome biosynthesis of *M. gryphiswaldense* (36), and was shown to 8-fold enhance the expression of a foreign gene compared to P_*mamDC*_ (35).

Indeed, mass spectra obtained by liquid chromatography – mass spectrometry (LC-MS) from extracts of strain *P_mamDC45_-trans-at-pks* showed a 7-fold increased intensity of the target mass, suggesting a successful transcriptional activation of the gene cluster (Fig. 2A). As the yield of the compound obtained from shake flasks cultures of *P_mamDC45_-trans-at-pks* was still too low for the isolation of the corresponding natural product, we scaled its production up to a 10L Fermenter, which provided enhanced aeration and growth of the culture. This approach enabled the isolation of 2 mg of the compound by semi-preparative HPLC, and the elucidation of its structure using MS and nuclear magnetic resonance spectrometry (NMR).

**Fig. 2.**
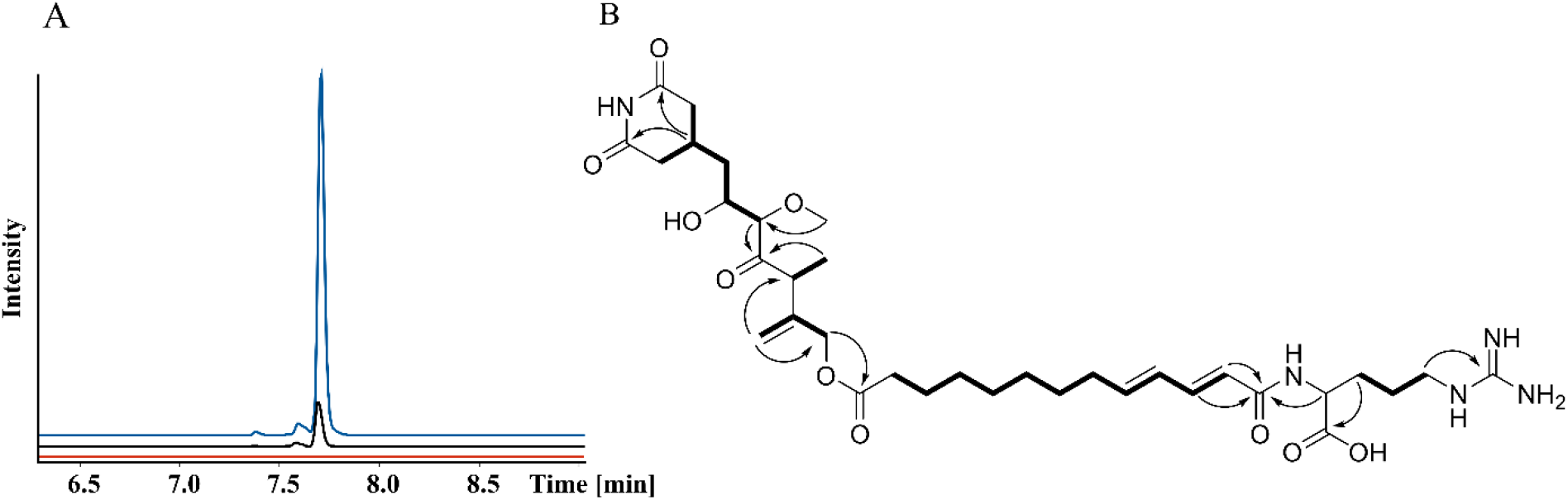
A: Extracted ion chromatograms for *m/z* 692.38 [M+H]^+^ showing the difference in compound production. In the strain Δ*trans-at-pks* (red), the production of the compound was abolished. In the promoter-activated strain *P_mamDC45_-trans-at-pks* (blue) the production was increased ca. 7-fold (AUC: 6513288) in comparison to the wild type (black) (AUC: 880064). B: NMR elucidated structure of sesbanimide R with the most relevant COSY and HMBC correlations.

### *De-novo* structure elucidation

HRESI-MS analysis of the compound (Fig. 2B) shows an [M+H]^+^ signal at *m/z* 692.3869 [M+H]^+^ (calc. 692.3865 Δ = −0.6 ppm) consistent with the neutral sum formula of C34H53N5O10 and containing 11 double bond equivalents (DBEs). The ^1^H-NMR and HSQC spectra of compound revealed a signal characteristic for an aliphatic exo double bond at *δ*(^1^H) = 5.18 and 4.96 ppm, which shows correlations to a methylene group at *δ*(^1^H) = 4.66 displayed as a singlet, as well as a methine group at *δ*(^1^H) = 3.71 ppm. The quartet of the methylene group shows COSY correlations to one methyl group at *δ*(^1^H) = 1.19 ppm and HMBC correlations to the quaternary carbon participating in the exo double bond at *δ*(^13^C) = 144.5 ppm besides a ketone at 213.2 ppm. The methylene group at *δ*(^1^H) = 4.66 therefore seems to be located between the ketone and the exo double bond, which is additionally confirmed by HMBC correlations of the methyl group at *δ*(^1^H) = 1.19 ppm to the ketone. On the other side of the ketone, another methine group with a chemical shift of *δ*(^1^H) = 3.66 ppm is found located based on their HMBC correlations. The high field chemical shift of this methine group in line with HMBC correlations to another methyl group with a high field chemical shift too at *δ*(^1^H) = 3.40 ppm reveals this part as methoxy function. The methine group at *δ*(^1^H) = 3.66 ppm furthermore shows COSY correlations to a second methine group at *δ*(^1^H) = 3.98 ppm. Its high field chemical shift suggests hydroxylation of this methine. It shows COSY correlations to a methylene group at *δ*(^1^H) = 1.49 ppm, which is located next to a methine group at 2.34 ppm based on their COSY correlations. This methine group exhibits further COSY correlations to two diastereotopic methylene groups at *δ*(^1^H) = 2.36, 2.68 and *δ*(^1^H) = 2.33, 2.70 ppm with almost identical chemical shifts, wherefore they have to be located in almost identical chemical surroundings. They do not reveal any further COSY, but HMBC correlations to two quaternary carbons at *δ*(^13^C) = 174.6 ppm. Based on the sum formula of the molecule and the 2D NMR data, the methine, the two methylene and the two quaternary carbons therefore likely are arranged as glutarimide, substituted in 4 position. There are no further correlations of any glutarimide participating functional groups, as a result this part depicts one end of the molecule.

Besides correlations of the methine group at *δ*(^1^H) = 4.66 ppm to the molecule part described above, it shows HMBC correlations to a quaternary carbon at *δ*(^13^C) = 14.6 ppm. Their high field chemical shifts suggest an ester bond in this position, which was confirmed by a saponification reaction (Fig. 3). The following seven methylene groups are arranged in a straight aliphatic chain, based on their chemical shifts and COSY as well as HMBC correlations. The high field chemical shift of the last of these seven methylene groups at *δ*(^1^H) = 2.18 ppm and its signals displayed as quartet suggest that it is followed by the first of four aromatic double bond protons at *δ*(^1^H) = 6.10, 6.22, 7.12 and 6.02 ppm. The two double bonds are conjugated based on COSY correlations of the four aromatic double bond protons and their high field carbon chemical shifts at *δ*(^13^C) = 144.1, 129.7, 142.3, 122.9 ppm. The last two double bond protons at *δ*(^1^H) = 7.12 and 6.02 ppm show HMBC correlations to a quaternary carbon at *δ*(^13^C) = 168.2 ppm. Its characteristic chemical shift and correlations of an alpha proton to this quaternary carbon reveal it as acid function of an amide bond. This alpha proton at *δ*(^1^H) = 4.40 ppm belongs to arginine, which was confirmed by Marfey’s analysis in addition to the following NMR correlations. It shows HMBC correlations to a free carboxylic acid function at *δ*(^13^C) = 177.6 ppm and two methylene groups at *δ*(^1^H) = 1.92 and 1.63 ppm, which themselves show HMBC correlations to a third more high field chemically shifted methylene group at *δ*(^1^H) = 3.22 ppm. Its characteristic chemical shift and correlations to a quaternary carbon at *δ*(^13^C) = 158.4 ppm reveals the coupling to the guanidine moiety of the molecule here, which marks the other end of compound. Additionally, to the NMR data we observed a fragment of *m/z* 397.245 [M+H]^+^ with LC-MS after saponification of the ester (Fig. 3). The size and sum formula of this fragment corresponds to the arginine containing part of the molecule and confirms the elucidated structure. This structure is highly similar to the structure of sesbanimide F, which became available at a late stage of our work in a study by Kacar et al. (28). However, our compound contains an additional terminal arginine (R) moiety. Hence, we used the name sesbanimide R.

**Fig. 3:**
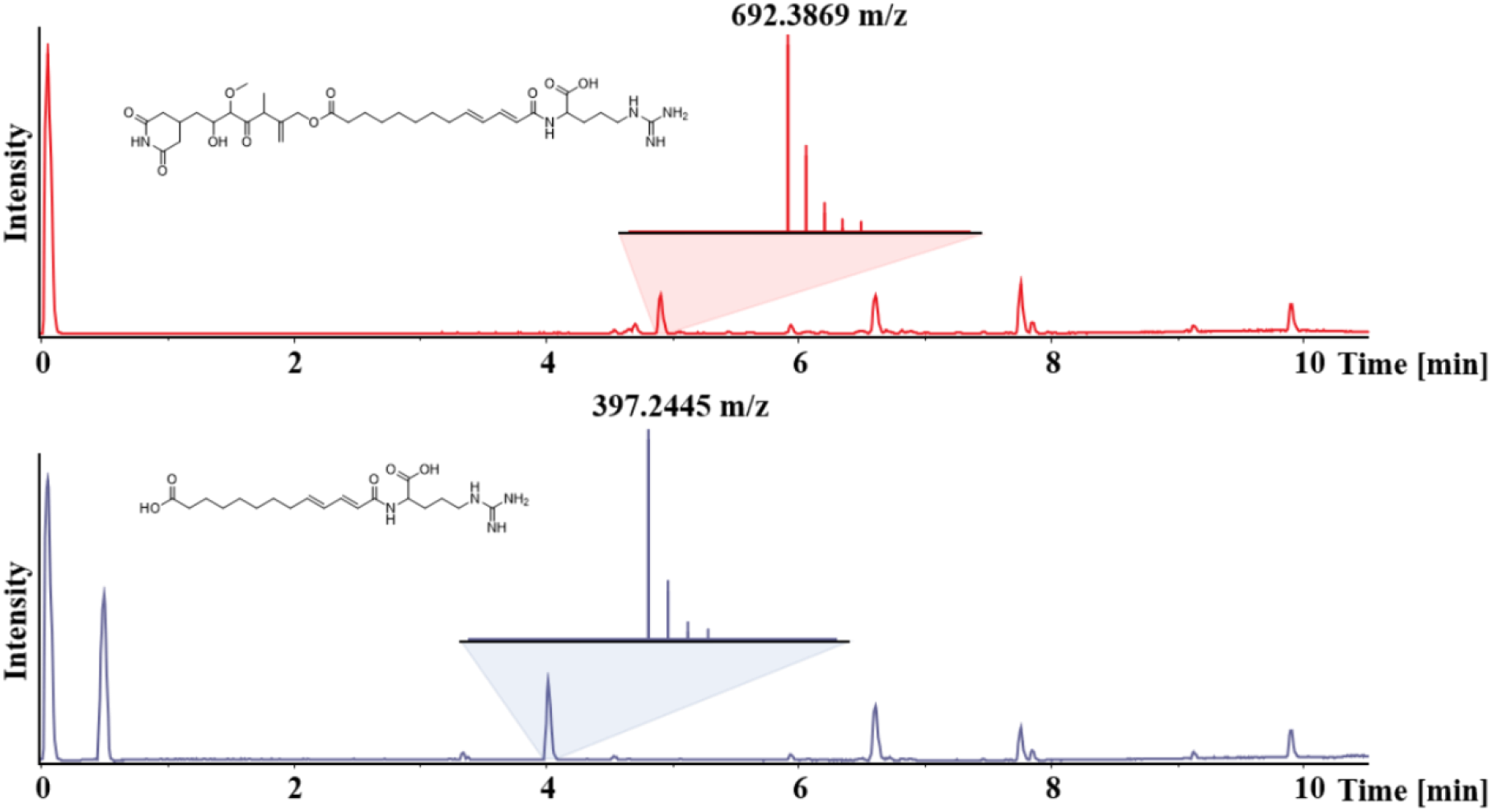
Saponification of Sesbanimide R to confirm the NMR elucidated structure. Base peak chromatogram of a sesbanimide R sample before (top) and after (bottom) treatment with NaOH. A fragment with an *m/z* of 397.2445 [M+H]^+^ was observed, which corresponds to the *arginine* containing part of the molecule after ester hydrolysis.

### Determination of the sesbanimide R stereochemistry

The vicinal coupling constant of 15.1 Hz for both aliphatic double bonds suggests an *E-*configuration of both double bonds. Marfey’s analysis and comparison to a commercially available L-Arginine standard revealed the Arginine from sesbanimide R to be S-configured. Due to instabilities of the molecule under acidic and basic conditions and selectivity issues between the free hydroxyl groups and the glutarimide, Mosher esterification experiments, which were carried out to elucidate the configuration of the remaining stereocenters, were not successful. When adding 10 or less equivalents of pyridine to the reaction mixture in chloroform, we observed complete degradation of the molecule. When performing the experiment in pure pyridine, the hydroxyl group undergoes fast elimination after formation of the respective Moshers ester. We were therefore not able to determine the absolute stereochemical configuration of the molecule experimentally and base our prediction of the stereochemistry on *in silico* analysis of the BGC. The transATor tool predicts the structure of *trans*-AT polyketides according to the substrate specificities of the involved KS domains (37). The top five hits of the tool predict the KS domain of module 4 to accept D-OH, while the sequence-based stereochemistry prediction for the KR-domain of module 3 was inconclusive. We therefore assign the stereocenter at C5 to be S-configured. Xie et al. (38) recently suggested that all C-methyl transferases in *trans*-AT PKS assembly lines generate (2R)-2-methyl-3-ketoacyl-ACP intermediates and that (2S)-2-methyl-3-hydroxyacyl-ACP intermediates are produced by epimerizing A2 type KR domains (38). As there is no KR domain present in module 4, we propose that the stereocenter at C8 is R-configured. The stereocenter at C6 is likely generated by a cyP450 enzyme (SbnE), but we were not able to make a prediction for its stereochemistry.

### *In silico* analysis of the gene cluster and biosynthesis hypothesis

A detailed annotation of the BGC was carried out (Table S2 in the Supporting Information). Besides using antiSMASH (30) for cluster and domain identification, additional information was gained by submitting the translated protein sequences to the TransATor tool (37) (Table S3 in the Supporting Information). Finally, the conserved domain search tool CDD was used to identify domains that were not identified by antiSMASH. The core biosynthetic gene cluster (BGC) spans over 39 kbp and consists of the two large pks genes *sbnO* (MSR-1_15630) and *sbnQ* (MSR-1_15650) as well as one monooxygenase encoding gene *sbnP* (MSR-1_15640). The core cluster is flanked by two AT domains encoded by *sbnA* and *sbnN.* SbnN was identified as an *in-trans* acyl transferase and SbnA as an *in-trans* acyl hydrolase. Several additional biosynthetic genes are encoded up and downstream of the core BGC: An asparagine synthase accompanied by an ACP domain (*sbnJ* and *sbnK*), a beta-branching cassette (41) (*sbnF-I*), a cytochrome p450 enzyme (*sbnE*), a methyltransferase (*sbnD*) and a standalone acyl-CoA dehydrogenase (*sbnX*). The ORFs 6, 8, 9, 11, 14 and 15 encode transport associated proteins putatively responsible for exporting sesbanimide R out of the cell. ORFs 2, 3, 5 and 7 encode regulatory proteins putatively responsible for controlling BGC expression and thereby sesbanimide production. ORFs 4, 10, 12 and 13 were annotated as encoding hypothetical proteins with unknown function. All KS domains of SbnO and SbnQ possess the actives site cysteine and histidine, except for the non-elongating KS5 of SbnQ which is missing the first histidine (42). All ACP domains of SbnO and SbnQ possess the canonical active site serine. The catalytic triad of serine, tyrosine and asparagine is present in all KR domains of SbnO and SbnQ. The DH1 of SbnO and DH1 and DH2 of SbnQ contain the conserved HxxxGxxxxP motive, which is missing in DH2 of SbnO and DH3 of SbnQ (Fig. S4 – S7 in the supporting information). We therefore propose the following biosynthesis scheme, based on *in silico* analysis of the BGC and considering the elucidated chemical structure of sesbanimide R (Fig. 4).

**Fig. 4:**
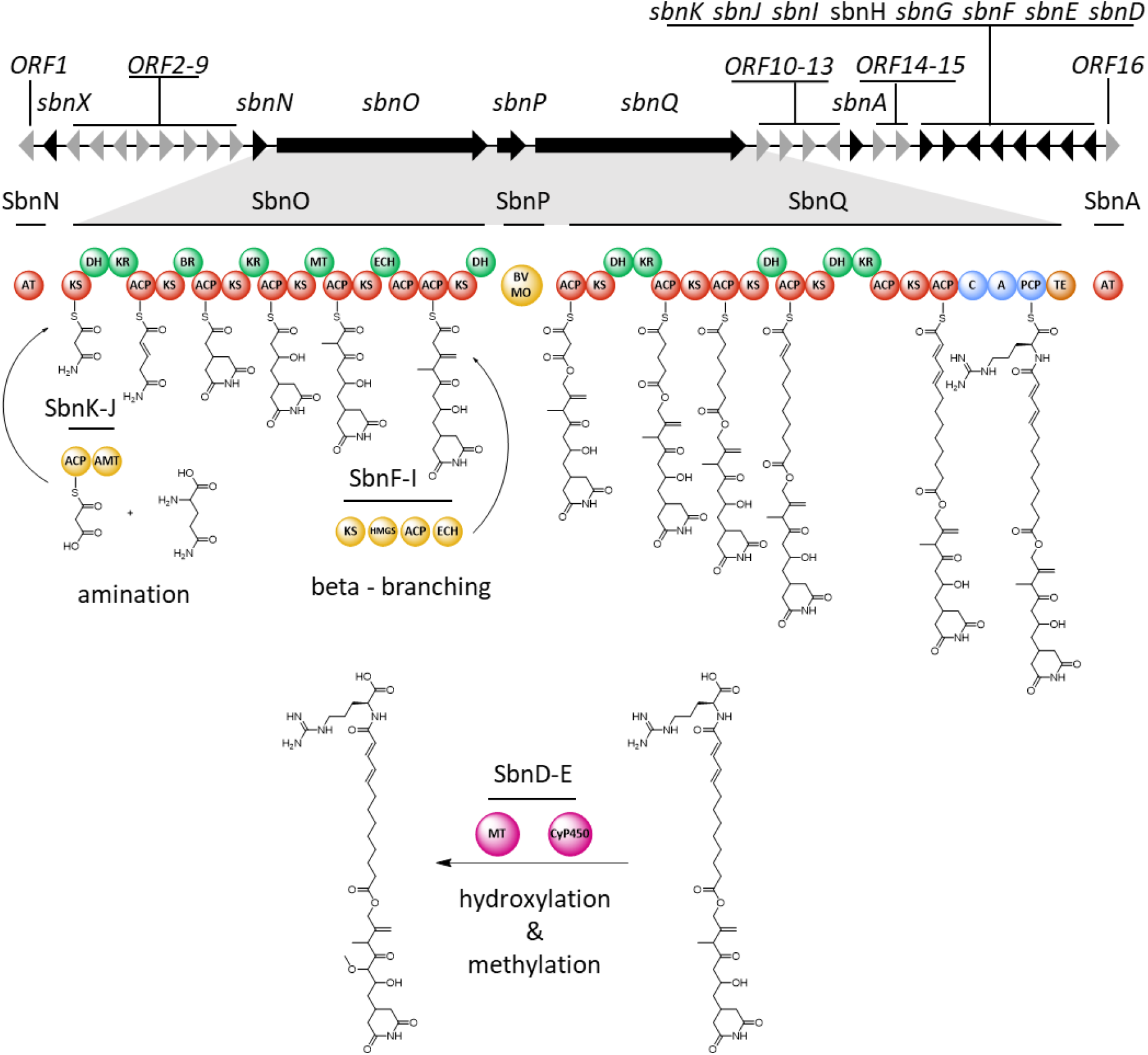
Proposed biosynthetic pathway for sesbanimide R. Core PKS modules are marked in red, core NRPS modules in blue and the thioesterase in orange. DH and KR modules of the core assembly line are marked in green. The amidotransferase, beta-branching cassette and Bayer-Villiger Monooxygenase are marked in yellow and the tailoring methyltransferase and Cyp450 enzyme in pink. The genes involved in the sesbanimide R biosynthesis are marked in black and named sbnA-X. The remaining genes with unknown or unassigned functioned are marked in grey and named ORF1-16

Initially, an amino group is transferred to ACP bound malonate by SbnJ (43). The starter moiety is then transferred to the first module of SbnO. Modules one and two of the assembly line then form the glutarimide moiety as previously described for the biosynthesis of Gladiofungin (44). Modules three to five elongate the nascent molecule according to the substrate specificity prediction for their KS domains. Exomethylene moiety incorporation by module five has previously been described in several *trans*-AT PKS biosyntheses (45). The domains required for exomethylene formation (ECH domain, tandem ACP domains and a beta branching cassette (*sbnF-I*)) are all present in the cluster. Module six is found split onto the genes *sbnO* and *sbnQ,* which are separated by *sbnP* encoding a Bayer-Villinger flavin-binding monooxygenase. Such a split module has also been shown to incorporate oxygen into polyketide backbones and is thus likely to be responsible for the ester formation in Sebanimide R (29). The second part of the molecule is synthesized by the PKS megasynthase SbnQ. Judging by the structure formula of sesbanimide R, we propose that module seven or eight performs one iterative PKS elongation step and thus incorporates a second malonyl-CoA building block into the final molecule. The DH and KR domains are proposed to act in *trans*, to biosynthesize the saturated part and double bonds present in sesbanimide R. An ER domain would also be needed to fully reduce the incorporated C2 unit, but this domain is not encoded on *sbnQ.* We therefore propose that this function is carried out by SbnX which was identified as an acyl-CoA dehydrogenase. The terminal NRPS module on *sbnQ* was predicted to incorporate L-arginine by NRPS predictor 2 which fits well to the elucidated structure (46). We propose that the methoxy group at C6 is incorporated by a cytochrome P450 enzyme and an Fkbm family methyltransferase, encoded by *sbnE* and *sbnD,* respectively.

Taken together, our devised biosynthesis scheme for sesbanimide R is very similar to the pathway suggested in parallel by Kacar et al. for the biosynthesis of sesbanimide F from *Stappia indica* PHM037 (28). The main differences between the two BGC lie in the distribution of DH and KR domains in SbnQ, the presence of three additional transport associated genes in the *M. gryphiswaldense* cluster, and a phosphopantetheinyl transferase in the *Stappia indica* cluster which is absent in the *M. gryphiswaldense* BGC. Notably, the final products from the strains under investigation by Kacar et al. do not contain the terminal arginine moiety observed in sesbanimide R even though the corresponding biosynthetic gene cluster contains the L-arginine-incorporating NRPS module (28). We speculate that the BGC from *M. gryphiswaldense* responsible for sesbanimide R formation is either an evolutionary intermediate in a developmental line leading to the sesbanimide gene cluster from PHM037 and PHM038, or that these clusters may carry a non-functional NRPS module. A conserved domain search of the A domains of the NRPS modules from *M. gryphiswaldense* and *S. indica* PHM037 revealed that the active sites are likely intact in both cases. In the case of the *S. indica* domain however, the residues just before the active site seem to be unusual for A domains. They were identified because they do not match the alignment against the reference A domains from the CDD database (Fig. S8 and S9 in the supporting information). Kačar et al. speculated that the arginine moiety is cleaved rapidly after the biosynthesis, so that the corresponding analogues are not detectable with the applied analytical conditions (28). As we were able to detect sesbanimide R, which was also relatively stable, we suggest as an alternative explanation that the uncommon residues close to the active site residues might result in an inactive A domain in the *S. indica* cluster, and that therefore no arginine is incorporated. Additionally, we did not detect any of the sesbanimides (A, B, C, D, E, and F) which were observed by Kacar et al. (28) in *M. gryphiswaldense*. A likely explanation might be that the tailoring steps resulting in the formation of the sesbanimides A, D, C and E only occur if no arginine moiety is present.

### Cytotoxicity

Sesbanimides have been associated with strong antitumor/cytotoxic activities which is common for polyketides containing a glutarimide moiety (48, 49). We therefore tested sesbanimide R *in vitro* against cell lines of liver carcinoma (HepG2), endocervical adenocarcinoma (KB3.1), colon carcinoma (HCT-116) and lung carcinoma (A549) (Table 1). The IC50 values against HePG2, HCT-116 and KB3.1 (1.4 – 4.3E-08 M) are comparable to sesbanimide F which exhibits a GI50 value of 2.0E-08 M against A549 cells (28). To better compare sesbanimide R and F, we also tested sesbanimide R against A549 cells which resulted in an IC50 of 3.0E-07. These results indicate that the arginine moiety either has no effect on cytotoxicity or reduces the cytotoxic effect in the case of A549 cells. The cytotoxicity of sesbanimide R falls well within the range commonly observed for glutarimide containing polyketides and other members of the sesbanimide compound family (28, 49, 50).

**Table 1:**
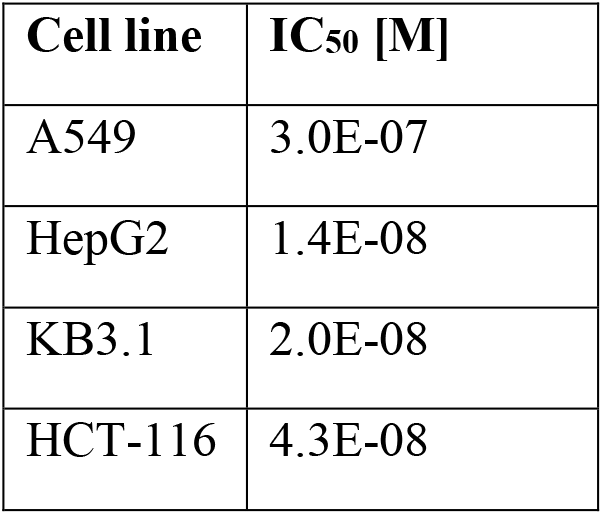
IC_50_ values of sesbanimide R against lung (A549), liver (HepG2), endocervical (KB3.1) and colon carcinoma (HCT-116).

| Cell line | IC<sub>50</sub> [M] |
| --- | --- |
| A549 | 3.0E-07 |
| HepG2 | 1.4E-08 |
| KB3.1 | 2.0E-08 |
| HCT-116 | 4.3E-08 |

## Conclusion

We unambiguously assigned a new member of the sesbanimide compound family to a *trans*-AT polyketide synthase biosynthetic gene cluster from *Magnetospirillum gryphiswaldense* by inactivation and overexpression of the cluster and statistical analysis of the strain’s metabolome.

Sesbanimide R belongs to the sesbanimide family of natural products. We suggest a biosynthesis pathway which is largely in line with the one proposed in a parallel study for sesbanimides A, C, D, E and F (28). In contrast to these compounds, sesbanimide R contains a terminal arginine moiety, which perfectly matches the *in silico* predictions of the BGC.

Sesbanimides were originally isolated from the seeds of *Sesbania drummondii* (51), and later from marine agrobacteria, indicating that symbiotic microorganisms are the actual sources for these metabolites rather than the plant (52, 28), a finding which is further supported by our study. Sesbanimide R is of interest owing to its cytotoxic bioactivity against several carcinoma cell lines, which is a characteristic of glutarimide containing polyketides (53, 49). The potent cytotoxic activity makes it a candidate for further investigations regarding its mode of action and development as an antitumor agent. As in other bacteria, the role of sesbanimide R for the physiology and fitness of *M. gryphiswaldense* in its freshwater habitat remains as yet elusive and requires further investigations.

Sesbanimide R is the first natural product identified and isolated from a magnetotactic bacterium. In addition to its well-established property to produce biogenic magnetic nanoparticles, it makes the tractable strain *M. gryphiswaldense* highly interesting also as a producer of secondary metabolites, another natural product of biotechnological relevance. Since numerous biosynthetic gene clusters encoding putative polyketide synthases and non-ribosomal peptide synthetases are present in the genomes of many different MTB (Table S4 in the Supporting Information), our study sets the stage for exploring this highly diverse group of prokaryotes as a potential source for the future discovery of novel secondary metabolites.

## Materials and Methods

### *In silico* analysis of the genome of magnetotactic bacteria and bioinformatics methods

The *Magnetospirillum gryphiswaldense* genome (accession no. CP027527) and genomes of other magnetotactic bacteria were screened for secondary metabolites biosynthetic gene clusters using the bioinformatic tool *antibiotics & Secondary Metabolite Analysis Shell* (antiSMASH version 5.1.2) (30). The amino acid sequence was aligned with the Basic Local Alignment Search Tool (BLASTp) against the publicly available database to find homologous proteins and to predict the functions of the ORFs. The presence of homologous ORFs in PHM037/PHM038 strains (28) was searched using the software Geneious Prime® (Biomatters Ltd., Auckland, New Zealand, 2020.0.3) (http://www.geneious.com). Furthermore, PKS and NRPS domain architecture and specificities present in the cluster were considered using TransAT or (http://transator.ethz.ch) and Pfam database (54).

### Bacterial strains and culture conditions

*Escherichia coli* was grown in lysogeny broth (LB) at 37°C and shaking at 180 rpm. Donor strain *E. coli* WM3064 (W. Metcalf, unpublished) was cultivated with 0.1 mM DL-α, f-diaminopimelic acid (DAP). *M. gryphiswaldense* was grown micro-aerobically at 28°C in modified flask standard medium (FSM) (33) with moderate agitation at 120 rpm, if not mentioned otherwise. Optical density (OD) and magnetic response (*C*_mag_) of *M. gryphiswaldense* strains were determined photometrically at 565 nm as reported earlier (31). Antibiotic selection was achieved by the addition of kanamycin at a concentration of 5 μg/ml (*M. gryphiswaldense*), and 25 μg/ml (*E. coli*) For Agar media, 1.5% (wt/vol) agar added to the liquid culture medium. Strains and vectors used in this study are given in Table S5.

### Molecular and genetic techniques

Oligonucleotides (Table S6) were purchased from Sigma-Aldrich (Steinheim, Germany). Chromosomal DNA of *M. gryphiswaldense* was isolated using a kit from Zymo Research, USA. Plasmids were constructed by standard recombinant techniques as described below. All constructs and selected amplicon from the mutants were sequenced by Macrogen Europe (Amsterdam, Netherlands).

### Construction of markerless site-specific deletion and activation of *trans*-AT PKS cluster

Markerless in-frame deletion of core-biosynthetic biosynthetic genes of *trans*-AT PKS cluster and insertion of a promoter in front of the cluster were conducted using homologous recombination-based on counter selection systems described previously (55). For the construction of the deletion plasmid, homologous regions of ca. 1.6 kb located upstream of MSR-1_15620 (Locus tag) including the first three codons of MSR-1_15620 and downstream of MSR-1_15650 with its last three codons were amplified from gDNA of *M. gryphiswaldense* using a proof-reading DNA polymerase and primer pairs RPA595/RPA596 and RPA597/RPA598. The PCR-products were purified from agarose gel using a gel extraction kit (Zymo Research, USA) and cloned into pORFM (55) digested with *Sal*I and *Not*I by Gibson assembly (56).

For activation of the *trans*-AT PKS cluster, the strong promoter *P_mamDC45_* with the spacing-optimized ribosome binding site (oRBS) was amplified from pAP150 (35) with primer pair RPA939/940. Homologous arms consisting of ca. 1.5 kb of C-terminus of MSR1_15590 and N-terminus of MSR1_15600 were amplified from gDNA of *M. gryphiswaldense* using primer pairs RPA937/938 and RPA940/941. The purified PCR products were assembled into pORFM (55), digested with *SalI* and *NotI* by Gibson assembly (56) with the *P_mamDC45_-oRBS* in between the two homologous arms. 5 μl of the Gibson assembly reaction was transformed into chemically competent *E. coli* DH5α (57), and the presence of the cloned fragment was confirmed by colony PCR using a pair of RPA484/485. The plasmid was isolated from the correct clone using Zymo Research kit, USA, and sequenced by Macrogen Europe (Armsterdam, Netherlands).

### Conjugation

Plasmid transfer by biparental conjugation was performed with donor strain *E. coli* WM3064 consisting of the verified construct and *M. gryphiswaldense* as the acceptor strain as reported previously (25). In-frame markerless chromosomal deletion and insertion were generated following the conjugative transfer of the plasmid to *M. gryphiswaldense* and homologous recombination utilizing GalK-based counterselection as previously described (55). Successful deletion and insertion yielded Δ*trans-at-pks* and *P_mamDC45_-trans-at-pks* strains, respectively. The mutants were confirmed by PCR using primers (Table S6) specific to sequences adjacent to the homologous regions, and verified by Sanger-sequencing of the amplicons.

### Growth curve and cell length analysis

For growth analyses, the strains were grown in 24 well plates (Sarstedt, Nümbrecht) in 1 ml FSM (33) in a microplate reader (infinite 200Pro, Tecan, Switzerland) with an automated reading of absorbance (560 nm) every 20 min for 150 cycles under aerobic conditions at 28°C shaking at 140 rpm. Absorbance values were corrected using FSM medium as blank. Cell length of the strains was estimated with the ImageJ plugin MicrobeJ 5.13i (58) using the ‘SHAPElength’ cell shape descriptor. Analysis of cell length was done as reported previously (59).

### Transmission electron microscopy

For TEM analysis, strains (wild type, Δ*trans-at-pks,* and *P_mamDC45_-trans-at-pks*) cultivated in 6 well plates (Sarstedt, Nümbrecht) under micro-oxic conditions at 24°C for 48 hrs, fixed in formaldehyde (1.8%), adsorbed onto carbon-coated copper grids (F200-CU Carbon support film, 200 Mesh, Electron Microscopy Sciences, Hatfield, UK), and washed three times with ddH2O. TEM was performed on JEM-2100 (JEOL Ltd., Tokyo, Japan) with an accelerating voltage of 80 kV. Images were captured with a Gatan Model 782 ES500W Erlangshen CCD camera (Gatan Inc., Pleasanton, USA) with the software Digital MicrographTM 1.80.70 (Gatan Inc., Pleasanton, USA). For data analysis and measurements, the software ImageJ Fiji V1.50c (60) was used.

### Screening for secondary metabolites and cultivation of strains

For the screening of secondary metabolites, *M. gryphiswaldense* and Δ*trans-at-pks* strains were cultivated at 28°C in FSM medium (33) with an initial OD565 of 0.01 under aerobic, microoxic and anaerobic conditions in 500 ml baffled Erlenmeyer flasks, and Duran Laboratory flasks with rubber-stopper containing 50 ml medium, and in 250ml Duran Laboratory flasks containing 240 ml degassed medium with rubber-stoppers, respectively. 1 mL (v/v) sterile amberlite resin XAD-16 (Sigma-Aldrich Chemie GmbH, Taufkirchen, Germany) was added to the culture grown under aerobic and micro-oxic conditions and 5 ml (v/v) XAD-16 into the culture grown under anaerobic conditions. The culture under aerobic condition was agitated at 150 rpm. The cells and the resin were harvested together by centrifugation after 60 hours of incubation before extraction.

To access the activation of the cluster, wild type, *P_mamDC45_-trans-at-pks,* and Δ*trans-at-pks* strains were cultivated under aerobic condition at 28°C in 100 ml FSM medium in 1L baffled Erlenmeyer flask with starting OD565 of 0.01 at 150 rpm. The culture was supplemented with 2 ml (v/v) sterile amberlite resin XAD-16 (Sigma-Aldrich Chemie GmbH, Taufkirchen, Germany). After 60 hours of incubation, the cells and resin were harvested together by centrifugation.

### Fermenter cultivation

Up-scale cultivation of the *P_mamDC45_-trans-at-pks* strain was done in a 10 L BioFlow® 120 Fermenter (Eppendorf) (34) under aerobic conditions with an initial OD565 of 0.04. A 900 ml culture grown under aerobic conditions was used as an inoculum for 9 L culture which was supplemented with 200 ml (v/v) XAD-16. The cells and resin were harvested together by centrifugation after 60 hours of incubation and dried in Lyophiliser before extraction.

### Extraction of the cell pellet and resin to screen for target masses

For screening of the target compound(s), the cell pellet and resin of each culture was extracted with 50 mL methanol for 1h. The extract was then dried and resolved in 2mL methanol. This extract was then centrifuged for 5 min at 215 g and diluted 1:10 prior to analysis with LC-MS system 1a and processing with metaboscape 5.0 (bruker).

### Extraction and isolation of sesbanimide R

The dry cells and resin from the upscaled fermentation were extracted three times with 500mL methanol. The extract was subsequently partitioned between hexane, ethylacetate and water. Sesbanimide R was detected solely in the aqueous layer. This layer was then dried and resuspended in methanol. Sesbanimide R was isolated from this pre-purified extract using LC-MS system 2. During purification, it became apparent, that sesbanimide R is unstable during prolonged exposure to light and oxygen simultaneously. Therefore, all purification steps were carried out with minimal exposure to light.

### LC-MS systems

All analytical LC-MS measurements were performed on a Dionex Ultimate 3000 RSLC system using a BEH C18, 100 x 2.1 mm, 1.7 μm dp column (Waters, Germany), coupled to a maXis 4G hr-ToF mass spectrometer (Bruker Daltonics, Germany) using the Apollo ESI source. UV spectra were recorded by a DAD in the range from 200 to 600 nm. The LC flow was split to 75 μL/min before entering the mass spectrometer.

#### LC-MS system 1a - standard measurements

Separation of 1 μl sample was achieved by a linear gradient from (A) H2O + 0.1 % FA to (B) ACN + 0.1 % FA at a flow rate of 600 μL/min and 45 °C. The gradient was initiated by a 0.5 min isocratic step at 5 % B, followed by an increase to 95 % B in 18 min to end up with a 2 min step at 95 % B before reequilibration under the initial conditions. Mass spectra were acquired in centroid mode ranging from 150 – 2500 *m/z* at a 2 Hz scan rate.

#### LC-MS system 1b – Marfey’s Method

Separation of 1 μl sample was achieved by a gradient from (A) H2O + 0.1 % FA to (B) ACN + 0.1 % FA at a flow rate of 600 μL/min and 45 °C. The gradient was as follows: Ramp in 1 min from 5% B to 10%B, in 14 min to 35% B, in 7 min to 55% B and in 3 min to 80 % B. This is followed by a 1 min step at 80% B before reequilibration with the initial conditions. Mass spectra were acquired in centroid mode ranging from 250 – 3000 *m/z* at a 2 Hz scan rate.

#### LC-MS system 1c – MS/MS measurements

Separation of 1 μl sample was achieved by a linear gradient from (A) H2O + 0.1 % FA to (B) ACN + 0.1 % FA at a flow rate of 600 μL/min and 45 °C. The gradient was initiated by a 0.5 min isocratic step at 5 % B, followed by an increase to 95 % B in 18 min to end up with a 2 min step at 95 % B before reequilibration under the initial conditions. Mass spectra were acquired in centroid mode ranging from 150 – 2500 *m/z* at a 2 Hz scan rate. Ions were selected for fragmentation by scheduled precursor list and the collision energy was determined by mass-and charge state dependent stepping from 25 to 60 eV.

#### LC-MS system 2

The final purification was performed on a Dionex Ultimate 3000 SDLC low pressure gradient system using a Luna, 5u, C18(2), 100A, 250 x 100 mm column (Phenomenex). Separation of 50 μl sample was achieved by a gradient from (A) H2O + 0.1 % FA to (B) ACN + 0.1 % FA at a flow rate of 5 mL/min and 45 °C. The gradient was as follows: A two min isocratic step at 5 %B, followed by a ramp to 35 %B in three min, ramp to 50 %B in 20 and to 95%B in 1 min. This 3 min wash step was followed by a return to initial conditions in 1 min and reequilibration for 3 min. UV spectra were recorded by a DAD in the range from 200 to 600 nm. The LC flow was split to 0.525 mL/min before entering the Thermo Fisher Scientific ISQ^TM^ EM single quadrupole mass spectrometer. Mass spectra were acquired by selected ion monitoring (SIM) at *m/z* 692.38 [M+H]^+^.

### Statistical analysis

Duplicates of wild type, and Δ*trans-at-pks* cultures were measured twice with LC-MS system 1a. Feature finding and bucketing was performed with the following parameters: Minimum Intensity: 5000; Minimum spectra for extraction: 5; minimum spectra for recursive feature extraction: 3. Recursive Feature extraction was performed when a feature was present in 2 out of 8 analyses and features were included in the bucket table when present in 3 out of 8 analyses after recursive feature extraction. Principal component analysis was performed to find differences between the two groups (wild type, and Δ*trans-at-pks*) with four analyses in each group. The PCA results were normalized with a logarithmic algorithm to account for low intensity features. Features that accounted for the largest difference between the data sets were reevaluated in the raw data.

### Marfey’s method to elucidate the stereochemistry of the arginine moiety

100 μg of sesbanimide R was dissolved in 100μL 6N HCL and incubated for 45 min at 110°C. It was subsequently dried under N2 stream and redisolved in 110μL dH2O. This was then split into 2 x 50 μL and 20 μL L-FDLA or D-FDLA and 20μL of NaHCO3 were added. The reaction was shaken at 700 rpm and 40°C for 2h and then stopped with the addition of 10μL 2N HCL. The reaction was then diluted with 300μL acetoneitrile, centrifuged and analyzed using LC-MS system 1b. The same reaction and measurement were performed with L-arginine as a reference.

### Saponification of sesbanimide R

50 μg of sesbanimide R were dried and redisolved in 100 μL 2 M NaOH. The reaction was stopped instantly by adding 200 μL1 M HCL and an aliquot of the solution was diluted 1:5 with acetonitrile and analyzed on LC-MS system 1a.

### Cytotoxicity assays with HCT-116, HepG2, KB3.1 cells and A549 cells

The cell lines were obtained from the German Collection of Microorganisms and Cell Cultures (Deutsche Sammlung für Mikroorganismen und Zellkulturen, DSMZ) and cultured under conditions recommended by the depositor. Cells were grown and diluted to 5 x 10^4^ cells per well of 96-well plates in 180 μL complete medium. After 2 h of equilibration, the cells were treated with a serial dilution of sesbanimide R in methanol. 20 μL of 5 mg/mL MTT (thiazolyl blue tetrazolium bromide) in PBS was added to each well after growing the cells for five days. The cells were further incubated for 2 h at 37°C, before the supernatant was discarded. Subsequently, the cells were washed with 100 μL PBS and treated with 100 μL 2-propanol/10 N HCl (250:1) to dissolve formazan granules. Cell viability was measured as percentage relative to the respective methanol control by measuring the absorbance at 570 nm with a microplate reader (Tecan Infinite M200Pro). Sigmoidal curve fitting was used to determine the IC50 values based on the absorbance measurements.

## Acknowledgements

This study was supported by the European Research Council (ERC) under the European Union’s Horizon 2020 research and innovation program (grant agreement no. 692637 to D.S.) and the Federal Ministry of Education and Research (BMBF) (Grant MagBioFab to D.S.). We thank Alexandra Amann and Stefanie Schmidt for performing the cytotoxicity assays and Daniel Krug and Fabian Panter for assistance with writing the manuscript.

## Author contributions

D.S., R.M., and R.P.A conceived and designed research; R.P.A, P.A.H., C.D. B., and C.N.R performed research; R.P.A., P.A.H., C.D.B, R.M., and D.S analyzed data; R.P.A., P.A.H., C.D.B., D.S., and R.M wrote the paper. All authors read and approved the final manuscript.

The authors declare no competing interest.

## References

1. Bazylinski DA, Lefèvre CT, Schüler D. 2013. Magnetotactic Bacteria, p. 453–494. In Rosenberg E, DeLong EF, Lory S, Stackebrandt E, Thompson F (ed), The Prokaryotes. Springer Berlin Heidelberg, Berlin, Heidelberg.

2. Uebe R, Schüler D. 2016. Magnetosome biogenesis in magnetotactic bacteria. Nat Rev Microbiol 14:621–637. doi:10.1038/nrmicro.2016.99.

3. McCausland HC, Komeili A. 2020. Magnetic genes: Studying the genetics of biomineralization in magnetotactic bacteria. PLoS Genet 16:e1008499. doi:10.1371/journal.pgen.1008499.

4. Lee J-H, Huh Y-M, Jun Y-w, Seo J-w, Jang J-t, Song H-T, Kim S, Cho E-J, Yoon H-G, Suh J-S, Cheon J. 2007. Artificially engineered magnetic nanoparticles for ultra-sensitive molecular imaging. Nat Med 13:95–99. doi:10.1038/nm1467.

5. Hergt R, Dutz S, Röder M. 2008. Effects of size distribution on hysteresis losses of magnetic nanoparticles for hyperthermia. J Phys Condens Matter 20:385214. doi:10.1088/0953-8984/20/38/385214.

6. Sun J, Li Y, Liang X-J, Wang PC. 2011. Bacterial Magnetosome: A Novel Biogenetic Magnetic Targeted Drug Carrier with Potential Multifunctions. J Nanomater 2011:469031–469043. doi:10.1155/2011/469031.

7. Mickoleit F, Lanzloth C, Schüler D. 2020. A Versatile Toolkit for Controllable and Highly Selective Multifunctionalization of Bacterial Magnetic Nanoparticles. Small 16:e1906922. doi:10.1002/smll.201906922.

8. Lefèvre CT, Bazylinski DA. 2013. Ecology, diversity, and evolution of magnetotactic bacteria. Microbiol Mol Biol Rev 77:497–526. doi:10.1128/MMBR.00021-13.

9. Lin W, Pan Y, Bazylinski DA. 2017. Diversity and ecology of and biomineralization by magnetotactic bacteria. Environ Microbiol Rep 9:345–356. doi:10.1111/1758-2229.12550.

10. Wenter R, Wanner G, Schüler D, Overmann J. 2009. Ultrastructure, tactic behaviour and potential for sulfate reduction of a novel multicellular magnetotactic prokaryote from North Sea sediments. Environ Microbiol 11:1493–1505. doi:10.1111/j.1462-2920.2009.01877.x.

11. Abreu F, Morillo V, Nascimento FF, Werneck C, Cantão ME, Ciapina LP, Almeida LGP de, Lefèvre CT, Bazylinski DA, Vasconcelos ATR de, Lins U. 2014. Deciphering unusual uncultured magnetotactic multicellular prokaryotes through genomics. ISME J 8:1055–1068. doi:10.1038/ismej.2013.203.

12. Monteil CL, Vallenet D, Menguy N, Benzerara K, Barbe V, Fouteau S, Cruaud C, Floriani M, Viollier E, Adryanczyk G, Leonhardt N, Faivre D, Pignol D, López-García P, Weld RJ, Lefevre CT. 2019. Ectosymbiotic bacteria at the origin of magnetoreception in a marine protist. Nat Microbiol 4:1088–1095. doi:10.1038/s41564-019-0432-7.

13. Lefèvre CT, Viloria N, Schmidt ML, Pósfai M, Frankel RB, Bazylinski DA. 2012. Novel magnetite-producing magnetotactic bacteria belonging to the Gammaproteobacteria. ISME J 6:440–450. doi:10.1038/ismej.2011.97.

14. Lin W, Zhang W, Zhao X, Roberts AP, Paterson GA, Bazylinski DA, Pan Y. 2018. Genomic expansion of magnetotactic bacteria reveals an early common origin of magnetotaxis with lineage-specific evolution. ISME J 12:1508–1519. doi:10.1038/s41396-018-0098-9.

15. Lin W, Zhang W, Paterson GA, Zhu Q, Zhao X, Knight R, Bazylinski DA, Roberts AP, Pan Y. 2020. Expanding magnetic organelle biogenesis in the domain Bacteria. Microbiome 8:152. doi:10.1186/s40168-020-00931-9.

16. Jogler C, Wanner G, Kolinko S, Niebler M, Amann R, Petersen N, Kube M, Reinhardt R, Schüler D. 2011. Conservation of proteobacterial magnetosome genes and structures in an uncultivated member of the deep-branching Nitrospira phylum. Proc Natl Acad Sci U S A 108:1134–1139. doi:10.1073/pnas.1012694108.

17. Kolinko S, Jogler C, Katzmann E, Wanner G, Peplies J, Schüler D. 2012. Single-cell analysis reveals a novel uncultivated magnetotactic bacterium within the candidate division OP3. Environ Microbiol 14:1709–1721. doi:10.1111/j.1462-2920.2011.02609.x.

18. Kolinko S, Wanner G, Katzmann E, Kiemer F, Fuchs BM, Schüler D. 2013. Clone libraries and single cell genome amplification reveal extended diversity of uncultivated magnetotactic bacteria from marine and freshwater environments. Environ Microbiol 15:1290–1301. doi:10.1111/1462-2920.12004.

19. Kolinko S, Richter M, Glöckner F-O, Brachmann A, Schüler D. 2014. Single-cell genomics reveals potential for magnetite and greigite biomineralization in an uncultivated multicellular magnetotactic prokaryote. Environ Microbiol Rep 6:524–531. doi:10.1111/1758-2229.12198.

20. Lefèvre CT, Trubitsyn D, Abreu F, Kolinko S, Jogler C, Almeida LGP de, Vasconcelos ATR de, Kube M, Reinhardt R, Lins U, Pignol D, Schüler D, Bazylinski DA, Ginet N. 2013. Comparative genomic analysis of magnetotactic bacteria from the Deltaproteobacteria provides new insights into magnetite and greigite magnetosome genes required for magnetotaxis. Environ Microbiol 15:2712–2735. doi:10.1111/1462-2920.12128.

21. Hoffmann T, Krug D, Bozkurt N, Duddela S, Jansen R, Garcia R, Gerth K, Steinmetz H, Müller R. 2018. Correlating chemical diversity with taxonomic distance for discovery of natural products in myxobacteria. Nat Commun 9:803. doi:10.1038/s41467-018-03184-1.

22. Araujo ACV, Abreu F, Silva KT, Bazylinski DA, Lins U. 2015. Magnetotactic bacteria as potential sources of bioproducts. Mar Drugs 13:389–430. doi:10.3390/md13010389.

23. Schleifer KH, Schüler D, Spring S, Weizenegger M, Amann R, Ludwig W, Köhler M. 1991. The Genus Magnetospirillum gen. nov. Description of Magnetospirillum gryphiswaldense sp. nov. and Transfer of Aquaspirillum magnetotacticum to Magnetospirillum magnetotacticum comb. nov. Systematic and Applied Microbiology 14:379–385. doi:10.1016/S0723-2020(11)80313-9.

24. Schüler D, Köhler M. 1992. The isolation of a new magnetic spirillum. Zentralblatt für Mikrobiologie 147:150–151. doi:10.1016/S0232-4393(11)80377-X.

25. Schultheiss D, Schüler D. 2003. Development of a genetic system for Magnetospirillum gryphiswaldense. Arch Microbiol 179:89–94. doi:10.1007/s00203-002-0498-z.

26. Zwiener Theresa, Dziuba Marina, Mickoleit Frank, Rückert Christian, Busche Tobias, Kalinowski Jörn, Uebe René, Schüler Dirk. Towards a ‘chassis’ for bacterial magnetosome biosynthesis: Genome streamlining of Magnetospirillum gryphiswaldense by multiple deletions. Microb Cell Fact.

27. Schüler D, Monteil CL, Lefevre CT. 2020. Magnetospirillum gryphiswaldense. Trends Microbiol 28:947–948. doi:10.1016/j.tim.2020.06.001.

28. Kačar D, Cañedo LM, Rodríguez P, Gonzalez E, Galán B, Schleissner C, Leopold-Messer S, Piel J, Cuevas C, La Calle F de, García JL. 2020. Identification of trans-AT polyketide clusters in two marine bacteria reveals cryptic similarities between distinct symbiosis factors. doi:10.1101/2020.09.18.303172.

29. Meoded RA, Ueoka R, Helfrich EJN, Jensen K, Magnus N, Piechulla B, Piel J. 2018. A Polyketide Synthase Component for Oxygen Insertion into Polyketide Backbones. Angew Chem Int Ed Engl 57:11644–11648. doi:10.1002/anie.201805363.

30. Blin K, Shaw S, Steinke K, Villebro R, Ziemert N, Lee SY, Medema MH, Weber T. 2019. antiSMASH 5.0: updates to the secondary metabolite genome mining pipeline. Nucleic Acids Res 47:W81–W87. doi:10.1093/nar/gkz310.

31. Schüler D, Uhl R, Bäuerlein E. 1995. A simple light scattering method to assay magnetism in Magnetospirillum gryphiswaldense. FEMS Microbiology Letters 132:139–145. doi:10.1111/j.1574-6968.1995.tb07823.x.

32. Cortina NS, Krug D, Plaza A, Revermann O, Müller R. 2012. Myxoprincomide: a natural product from Myxococcus xanthus discovered by comprehensive analysis of the secondary metabolome. Angew Chem Int Ed Engl 51:811–816. doi:10.1002/anie.201106305.

33. Heyen U, Schüler D. 2003. Growth and magnetosome formation by microaerophilic Magnetospirillum strains in an oxygen-controlled fermentor. Appl Microbiol Biotechnol 61:536–544. doi:10.1007/s00253-002-1219-x.

34. Riese CN, Uebe R, Rosenfeldt S, Schenk AS, Jérôme V, Freitag R, Schüler D. 2020. An automated oxystat fermentation regime for microoxic cultivation of Magnetospirillum gryphiswaldense. Microb Cell Fact 19:206. doi:10.1186/s12934-020-01469-z.

35. Borg S, Hofmann J, Pollithy A, Lang C, Schüler D. 2014. New vectors for chromosomal integration enable high-level constitutive or inducible magnetosome expression of fusion proteins in Magnetospirillum gryphiswaldense. Appl Environ Microbiol 80:2609–2616. doi:10.1128/AEM.00192-14.

36. Schübbe S, Würdemann C, Peplies J, Heyen U, Wawer C, Glöckner FO, Schüler D. 2006. Transcriptional organization and regulation of magnetosome operons in Magnetospirillum gryphiswaldense. Appl Environ Microbiol 72:5757–5765. doi:10.1128/AEM.00201-06.

37. Helfrich EJN, Ueoka R, Dolev A, Rust M, Meoded RA, Bhushan A, Califano G, Costa R, Gugger M, Steinbeck C, Moreno P, Piel J. 2019. Automated structure prediction of trans-acyltransferase polyketide synthase products. Nat Chem Biol 15:813–821. doi:10.1038/s41589-019-0313-7.

38. Xie X, Khosla C, Cane DE. 2017. Elucidation of the Stereospecificity of C-Methyltransferases from trans-AT Polyketide Synthases. J. Am. Chem. Soc. 139:6102–6105. doi:10.1021/jacs.7b02911.

39. Blin K, Shaw S, Kautsar SA, Medema MH, Weber T. 2020. The antiSMASH database version 3: increased taxonomic coverage and new query features for modular enzymes. Nucleic Acids Res. doi:10.1093/nar/gkaa978.

40. Lu S, Wang J, Chitsaz F, Derbyshire MK, Geer RC, Gonzales NR, Gwadz M, Hurwitz DI, Marchler GH, Song JS, Thanki N, Yamashita RA, Yang M, Zhang D, Zheng C, Lanczycki CJ, Marchler-Bauer A. 2020. CDD/SPARCLE: the conserved domain database in 2020. Nucleic Acids Res 48:D265–D268. doi:10.1093/nar/gkz991.

41. Kjaerulff L, Raju R, Panter F, Scheid U, Garcia R, Herrmann J, Müller R. 2017. Pyxipyrrolones: Structure Elucidation and Biosynthesis of Cytotoxic Myxobacterial Metabolites. Angew Chem Int Ed Engl 56:9614–9618. doi:10.1002/anie.201704790.

42. Nguyen T, Ishida K, Jenke-Kodama H, Dittmann E, Gurgui C, Hochmuth T, Taudien S, Platzer M, Hertweck C, Piel J. 2008. Exploiting the mosaic structure of trans-acyltransferase polyketide synthases for natural product discovery and pathway dissection. Nat Biotechnol 26:225–233. doi:10.1038/nbt1379.

43. Lešnik U, Lukežič T, Podgoršek A, Horvat J, Polak T, Šala M, Jenko B, Harmrolfs K, Ocampo-Sosa A, Martínez-Martínez L, Herron PR, Fujs Š, Kosec G, Hunter IS, Müller R, Petković H. 2015. Construction of a new class of tetracycline lead structures with potent antibacterial activity through biosynthetic engineering. Angew Chem Int Ed Engl 54:3937–3940. doi:10.1002/anie.201411028.

44. Niehs SP, Kumpfmüller J, Dose B, Little RF, Ishida K, Flórez LV, Kaltenpoth M, Hertweck C. 2020. Insect-Associated Bacteria Assemble the Antifungal Butenolide Gladiofungin by Non-Canonical Polyketide Chain Termination. Angew Chem Int Ed Engl 59:23122–23126. doi:10.1002/anie.202005711.

45. Helfrich EJN, Piel J. 2016. Biosynthesis of polyketides by trans-AT polyketide synthases. Nat Prod Rep 33:231–316. doi:10.1039/c5np00125k.

46. Röttig M, Medema MH, Blin K, Weber T, Rausch C, Kohlbacher O. 2011. NRPSpredictor2—a web server for predicting NRPS adenylation domain specificity. Nucleic Acids Res. 39:W362–W367. doi:10.1093/nar/gkr323.

47. Bhushan R, Bruckner H. 2004. Marfey’s reagent for chiral amino acid analysis: A review. Amino Acids 27:231–247. doi:10.1007/s00726-004-0118-0.

48. Gorst-Allman CP, Steyn PS, Vleggaar R, Grobbelaar N. 1984. Structure elucidation of sesbanimide using high-field n.m.r. spectroscopy. J Chem Soc Perk T 1:1311. doi:10.1039/p19840001311.

49. Rajski SR, Shen B. 2010. Multifaceted modes of action for the glutarimide-containing polyketides revealed. Chembiochem 11:1951–1954. doi:10.1002/cbic.201000370.

50. Sugawara K, Nishiyama Y, Toda S, Komiyama N, Hatori M, Moriyama T, Sawada Y, Kamei H, Konishi M, Oki T. 1992. Lactimidomycin, a new glutarimide group antibiotic. Production, isolation, structure and biological activity. J Antibiot (Tokyo) 45:1433–1441. doi:10.7164/antibiotics.45.1433.

51. Powell RG, Smith CR, Weisleder D, Matsumoto G, Clardy J, Kozlowski J. 1983. Sesbanimide, a potent antitumor substance from Sesbania drummondii seed. J. Am. Chem. Soc. 105:3739–3741. doi:10.1021/ja00349a081.

52. Acebal C, Alcazar R, Cañedo LM, La Calle F de, Rodriguez P, Romero F, Fernandez Puentes JL. 1998. Two marine Agrobacterium producers of sesbanimide antibiotics. J Antibiot (Tokyo) 51:64–67. doi:10.7164/antibiotics.51.64.

53. Gorst-Allman CP, Steyn PS, Vleggaar R, Grobbelaar N. 1984. Structure elucidation of sesbanimide using high-field n.m.r. spectroscopy. J. Chem. Soc., Perkin Trans. 1:1311. doi:10.1039/P19840001311.

54. Finn RD, Coggill P, Eberhardt RY, Eddy SR, Mistry J, Mitchell AL, Potter SC, Punta M, Qureshi M, Sangrador-Vegas A, Salazar GA, Tate J, Bateman A. 2016. The Pfam protein families database: towards a more sustainable future. Nucleic Acids Res 44:D279–85. doi:10.1093/nar/gkv1344.

55. Raschdorf O, Plitzko JM, Schüler D, Müller FD. 2014. A tailored galK counterselection system for efficient markerless gene deletion and chromosomal tagging in Magnetospirillum gryphiswaldense. Appl Environ Microbiol 80:4323–4330. doi:10.1128/AEM.00588-14.

56. Gibson DG, Young L, Chuang R-Y, Venter JC, Hutchison CA, Smith HO. 2009. Enzymatic assembly of DNA molecules up to several hundred kilobases. Nat Methods 6:343–345. doi:10.1038/nmeth.1318.

57. Hanahan D. 1983. Studies on transformation of Escherichia coli with plasmids. Journal of Molecular Biology 166:557–580. doi:10.1016/s0022-2836(83)80284-8.

58. Ducret A, Quardokus EM, Brun YV. 2016. MicrobeJ, a tool for high throughput bacterial cell detection and quantitative analysis. Nat Microbiol 1:16077. doi:10.1038/nmicrobiol.2016.77.

59. Pfeiffer D, Toro-Nahuelpan M, Awal RP, Müller F-D, Bramkamp M, Plitzko JM, Schüler D. 2020. A bacterial cytolinker couples positioning of magnetic organelles to cell shape control. Proc Natl Acad Sci U S A. doi:10.1073/pnas.2014659117.

60. Schindelin J, Arganda-Carreras I, Frise E, Kaynig V, Longair M, Pietzsch T, Preibisch S, Rueden C, Saalfeld S, Schmid B, Tinevez J-Y, White DJ, Hartenstein V, Eliceiri K, Tomancak P, Cardona A. 2012. Fiji: an open-source platform for biological-image analysis. Nat Methods 9:676–682. doi:10.1038/nmeth.2019.

