## Supplementary Data for "Sesbanimide R, a novel cytotoxic polyketide produced by magnetotactic bacteria"

Table S1: Putative secondary metabolites gene clusters present in the genome of *Magnetospirillum gryphiswaldense* with locus tag and size.

| Gene Clusters | Locus tags |
|---|---|
| Lasso peptide | MSR-1_06390-06890 |
| Aryl polyene | MSR-1_09890-10640 |
| Homoserine lactone | MSR-1_16040-16260 |
| Trans-AT PKS | MSR-1_15520-15810 |

**Schemes for the inactivation of the *trans*-AT PKS gene cluster from *Magnetospirillum gryphiswaldense***

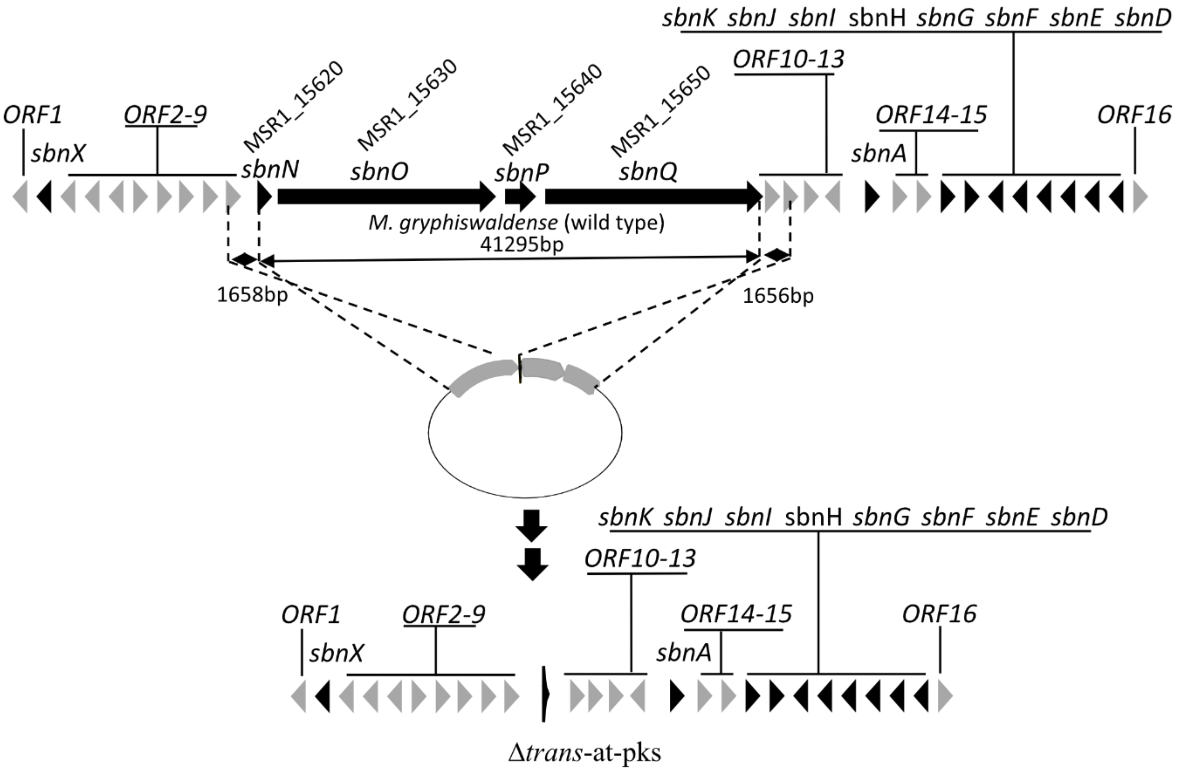


Fig. S1: Simplified schematic illustration of an in-frame deletion of core-biosynthetic genes of *trans*-AT PKS cluster in *Magnetospirillum gryphiswaldense,* yielding strain Δ*trans-at-pks*.

**Identification of sesbanimide R as the target mass by principal component analysis:**

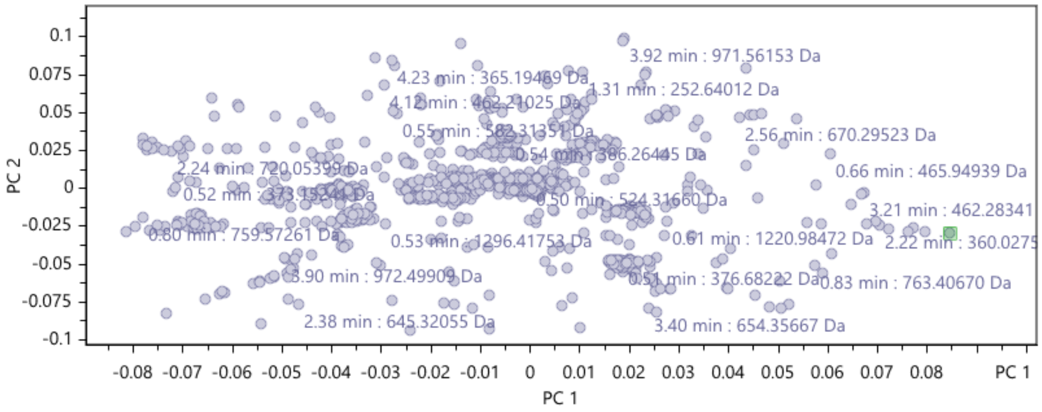


Fig. S2: Principal component analysis of the extracts of the *M. gryphiswaldense* wildtype and Δ*trans-at-pks* strain. The more a component is responsible for a difference between the data sets, the farther it is to the right (wildtype) or to the left (Δ*trans-at-pks*) from the center, the highlighted feature represents the target mass 692.38 m/z which was assigned to the trans-AT PKS cluster and was found only in the wildtype strain.

**Schemes for the activation of the *trans*-AT PKS gene cluster from *Magnetospirillum gryphiswaldense***

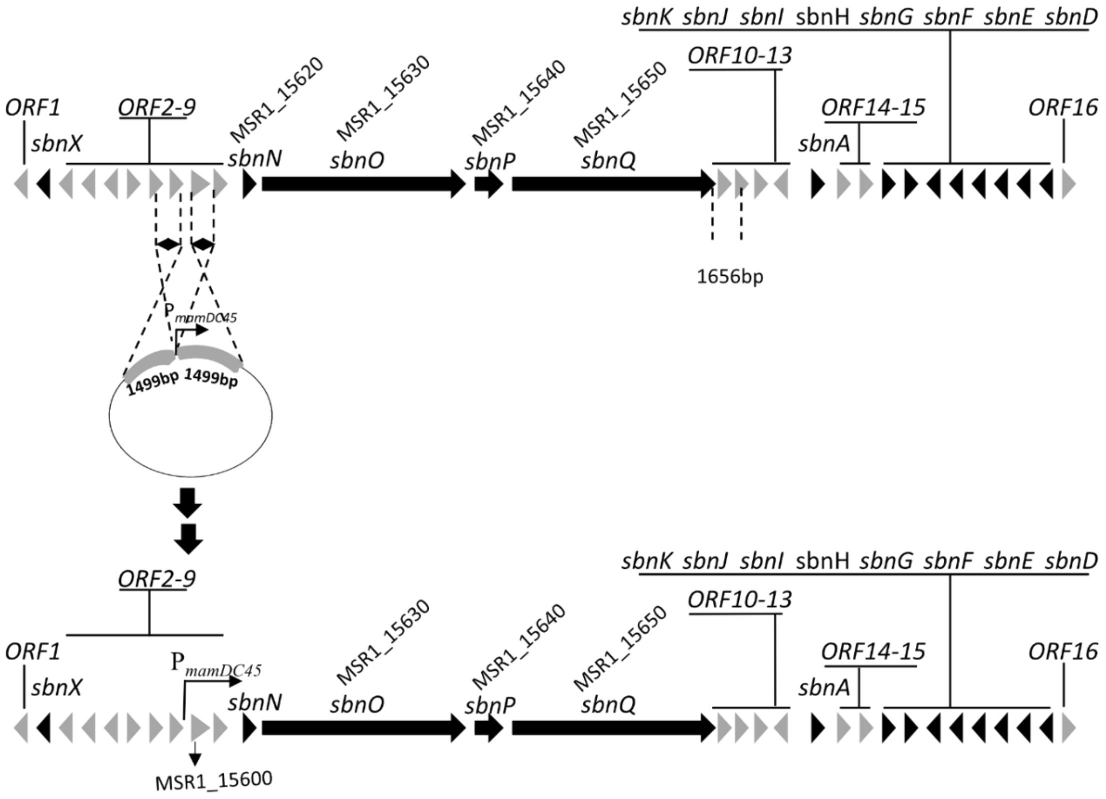


Fig. S3: Simplified schematic illustration of the insertion of promoter $P_{mamDC45}$ in front of MSR-1_15600 in *trans*-AT PKS cluster in *Magnetospirillum gryphiswaldense,* generating strain $P_{mamDC45}$*-trans-at-pks*.

### In silico analysis of the sesbanimide biosynthetic gene cluster from *Magnetospirillum gryphiswaldense*

Table S2: Annotation of the sesbanimide gene cluster in *Magnetospirillum gryphiswaldense* and comparison the sesbanimide cluster in PHM037 and PHM038

| | **Putative function/ homologue** | **Accession number of closest protein homologue** | **Cover/Pairwise identity [%]** | **Corresponding gene in PHM037/ PHM038** |
|---|---|---|---|---|
| Orf1 | CoA transferase | WP_024079852 | 100/100 | - |
| Orf2 | response regulator | WP_024079854 | 100/100 | - |
| Orf3 | response regulator receiver modulated diguanylate cyclase | CDK98834 | 100/100 | - |
| Orf4 | Hpt domain-containing protein | WP_024079856 | 73.5/100 | - |
| Orf5 | diguanylate cyclase | WP_024079857 | 90.4/100 | - |
| Orf6 | amino acid ABC transporter substrate-binding protein | WP_024079858 | 100/100 | - |
| Orf7 | response regulator | WP_024079859 | 100/100 | - |
| Orf8 | cyclic peptide export ABC transporter | WP_024079860 | 100/100 | SbnL |
| Orf9 | cyclic peptide export ABC transporter | WP_024079861 | 100/100 | SbnM |
| Orf10 | Band 7 protein | CDK98845 | 100/100 | SbnR |
| Orf11 | ABC transporter substrate-binding protein | WP_024079867 | 100/100 | SbnS |
| Orf12 | hypothetical protein | WP_024079868 | 100/100 | SbnT |
| Orf13 | DUF-697 domain-containing protein | WP_024079869 | 94.7/100 | SbnU |
| Orf14 | putative lysine-arginine-ornithine-binding periplasmic protein | CDK98850 | 100/100 | - |
| Orf15 | cation:dicarboxylase symporter family transporter | WP_024079872 | 88.7/99.8 | - |
| Orf16 | metallophosphoesterase | WP_041633527 | 100/100 | SbnC |
| SbnA | acyltransferase domain-containing protein | WP_158497748 | 73.8/99.7 | SbnA |
| SbnD | Fkbm family methyltransferase | WP_158497749 | 87.2/100 | SbnD |
| SbnE | cytochrome P450 | WP_024079879 | 100/100 | SbnE |
| SbnF | hydroxymethylglutaryl-CoA synthase family protein | WP_024079878 | 100/100 | SbnF |
| SbnG | acyl carrier protein | WP_024079877 | 100/100 | SbnG |
| SbnH | decarboxylase | WP_024079876 | 100/100 | SbnH |
| SbnI | enoyl-CoA hydratase/ isomerase | WP_024079875 | 100/100 | SbnI |
| SbnJ | asparagine synthase (glutamine-hydrolyzing) | WP_024079874 | 100/100 | SbnJ |
| SbnK | acyl carrier protein | WP_106002060 | 100/96.3 | SbnK |
| SbnN | ACP S-malonyltransferase | WP_024079862 | 100/100 | SbnN |
| SbnO | SDR family NAD(P)-dependent oxidoreductase | WP_024079863 | 100/100 | SbnO |
| SbnP | monooxygenase | OJX77727 | 99.5/87.0 | SbnP |
| SbnQ | non-ribosomal peptide synthetase | WP_024079865 | 100/100 | SbnQ |
| SbnX | acyl-CoA dehydrogenase | WP_024079853 | 100/100 | SbnX |

Table S3: Substrate specificities as predicted by the TranATor tool and sorted according to E-value. The predictions that fit the structure and biosynthesis proposal are highlighted in bold.

| | **prediction** | | **e value** | **score** |
|---|---|---|---|---|
| KS1 (sbnO) | Clade 95 | various specificities | 1.1E-179 | 590.7 |
| | Clade 109 | completely reduced | 1.0E-176 | 580.7 |
| | **Clade 8** | **unusual starter (AMT/Succinate)** | 2.8E-217 | 714.5 |
| | Clade 136 | β D-OH | 2.8E-178 | 585.8 |
| | Clade 96 | various specificies (mainly α-Me) | 2.0E-177 | 583.2 |
| KS2 (sbnO) | Clade_64 | non-elongating (double bonds (mostly z-configured)) | 9.1E-218 | 715.9 |
| | Clade 5 | amino acids (oxa/thia) | 2.9E-151 | 497.0 |
| | **Clade 82** | **double bonds (mostly e-configured)** | 5.8E-151 | 495.9 |
| | **Clade 115** | **β-keto or double bonds** | 7.8E-149 | 488.8 |
| | **Clade 99** | **double bonds (e-configured)** | 4.4E-147 | 483.4 |
| KS3 (sbnO) | Clade 95 | various specificities | 2.1E-149 | 490.9 |
| | Clade 109 | completely reduced | 1.4E-141 | 464.9 |
| | Clade 96 | various specificies (mainly α-Me) | 4.3E-166 | 545.8 |
| | **Clade 12** | **vinylogous chain branching** | 2.5E-155 | 510.2 |
| | Clade 136 | β D-OH | 5.0E-153 | 502.6 |
| KS4 (sbnO) | **Clade_7** | **ß D-OH** | 1.2E-207 | 683.1 |
| | **Clade_110** | **ß D-OH or double bonds (e-configured)** | 8.7E-212 | 696.5 |
| | **Clade_140** | **β D-OH** | 9.2E-209 | 686.7 |
| | **Clade_137** | **β D-OH** | 8.9E-207 | 680.0 |
| | **Clade_62** | **β D-OH (some with α L-Me)** | 1.6E-201 | 662.6 |
| KS5 (sbnO) | Clade_68 | α L-OH/Me β D-OH | 8.6E-174 | 571.1 |
| | Clade_104 | ß OMe or ß Me double bond | 8.4E-170 | 557.7 |
| | **Clade_21** | **α Me reduced/keto/D-OH** | 2.5E-194 | 638.7 |
| | **Clade_74** | **α Me reduced/keto/D-OH** | 7.3E-193 | 633.9 |
| | **Clade_23** | **α-Me** | 5.9E-180 | 591.3 |
| KS6 (sbnO) | Clade_104 | ß OMe or ß Me double bond | 1.9E-177 | 582.9 |
| | Clade_86 | α-L-Me red or OH | 6.7E-169 | 554.9 |
| | Clade_14 | exomethyl/exoester | 9.3E-196 | 643.5 |
| | Clade_2 | α-Me shifted double bond or OH | 3.6E-174 | 572.2 |
| | **Clade_73** | **exomethylene** | 1.1E-169 | 557.3 |
| KS7 (sbnQ) | **Clade_35** | **oxidative rearrangement** | 4.7E-214 | 704.0 |
| | Clade_95 | various specificities | 9.4E-167 | 548.0 |
| | Clade_25 | completely reduced | 8.3E-173 | 567.8 |
| | Clade_96 | various specificies (mainly α-Me) | 3.7E-172 | 565.8 |
| | Clade_136 | β D-OH | 8.9E-169 | 554.5 |
| KS8 (sbnQ) | Clade_95 | various specificities | 2.6E-181 | 596.0 |
| | **Clade_25** | **completely reduced** | 1.2E-201 | 662.9 |
| | Clade_108 | shifted double bonds | 5.2E-191 | 627.9 |
| | Clade_96 | various specificies (mainly α-Me) | 9.3E-188 | 617.2 |
| | Clade_136 | β D-OH | 2.2E-173 | 569.7 |
| KS9 (sbnQ) | **Clade_25** | **completely reduced** | 9.4E-217 | 712.6 |
| | Clade_108 | shifted double bonds | 2.9E-204 | 671.6 |
| | Clade_96 | various specificies (mainly α-Me) | 1.2E-190 | 626.7 |

| | | | | |
|---|---|---|---|---|
| | Clade_11 | shifted double bonds | 6.5E-189 | 621.0 |
| | Clade_136 | β D-OH | 5.4E-185 | 607.9 |
| KS10 (sbnQ) | **Clade_82** | **double bonds (mostly e-configured)** | 1.5E-227 | 748.3 |
| | **Clade_125** | **double bonds (e-configured)** | 6.4E-223 | 733.0 |
| | **Clade_129** | **double bonds (e-configured)** | 6.5E-213 | 700.1 |
| | **Clade_101** | **double bonds** | 9.2E-213 | 699.5 |
| | **Clade_99** | **double bonds (e-configured)** | 9.9E-213 | 699.7 |
| KS11 (sbnQ) | **Clade_76** | **non-elongating (double bonds)** | 1.7E-179 | 589.9 |
| | **Clade_142** | **non-elongating (various)** | 2.7E-167 | 549.6 |
| | Clade_101 | double bonds | 1.6E-162 | 534.0 |
| | Clade_90 | β-keto or double bonds | 4.1E-161 | 529.5 |
| | Clade_115 | β-keto or double bonds | 8.7E-160 | 524.9 |

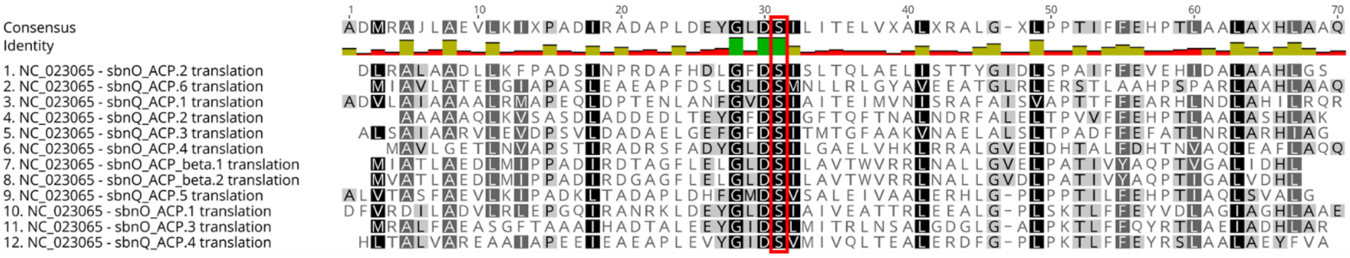


Fig. S4: Aligment of ACP domains of *sbnO* and *sbnQ*. All ACPs contain the active site serine.

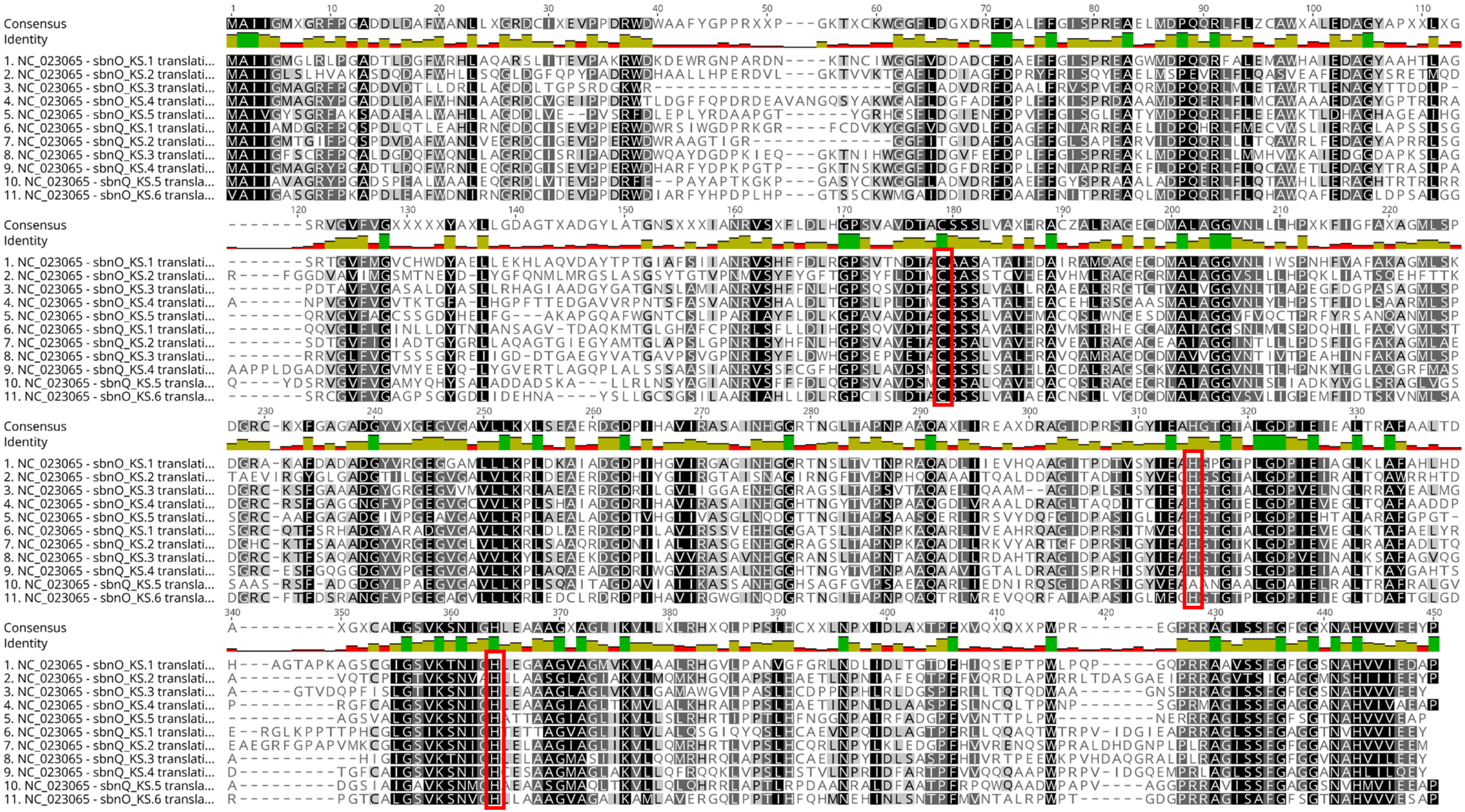

Fig. S5: Alignment of KS domains of *sbnO* and *sbnQ*. All KS domains contain the conserved cysteine and two histidines except for KS5 of *sbnQ*

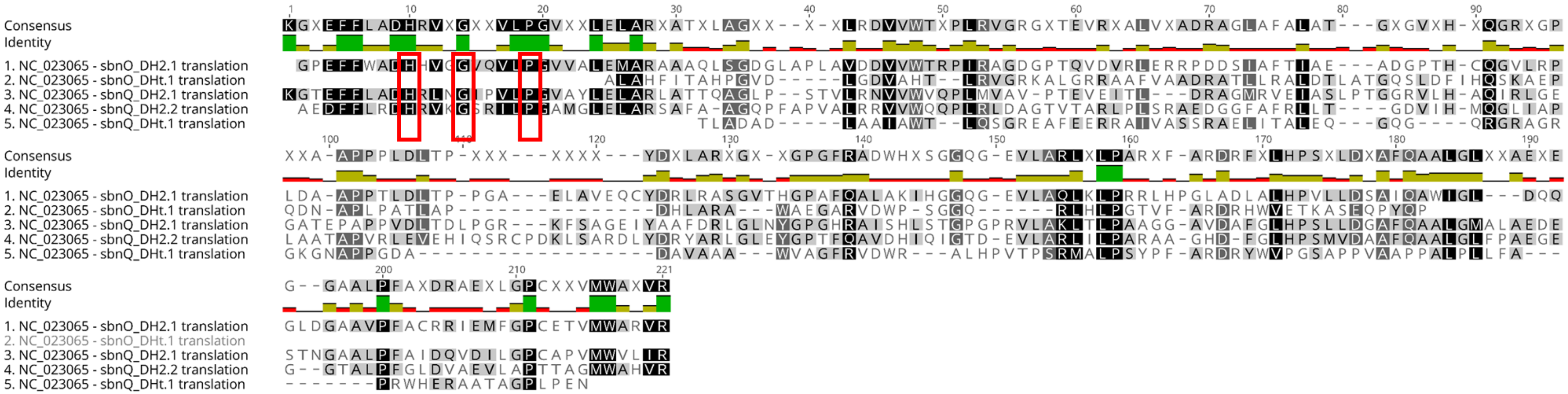

Figure S6. Alignment of DH domains of *sbnO* and *sbnQ*. DHt1 of *sbnO* and DH1 of *sbnQ* are missing the conserved HxxxGxxxxP motive.

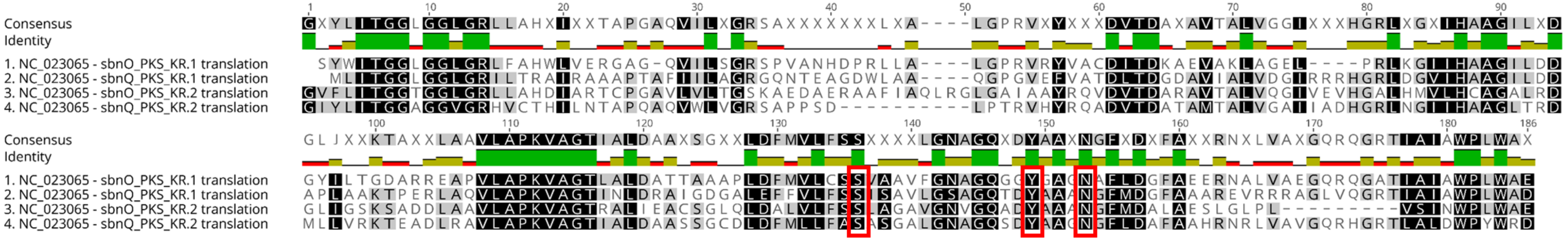

Fig. S7: Alignment of KR domains of *sbnO* and *sbnQ*. All domains contain the conserved active site residues serine, tyrosine and asparagine.

```
Feature 2
query          16 GETLTARQLLRQVERLCLILSSCG.[4].DRVAVL.[3].EAGGIAGLLSVLASGAVWVPLDPEHP.[1].ERLSLILRD 88
1AMU_A         62 NEQLTYHELNVKANQLARIFIEKG.[4].TLVGIM.[3].SIDLFIGILAVLKAGGAYVPIDIEYP.[1].ERIQYILDD 134  Brevibacillus brevis
3VNQ_A         58 DDRISYGRLDAWSDAVARTLLAEG.[4].DRVALR.[3].GAEAIVAILAILKCGAAYVPVDLRNP.[1].SRSDFILAD 130  Streptomyces sp.
4ZXH_A        504 DHTVSYRELGQHIAGIAEYLRAHG.[4].DRVGLM.[3].TALLPAAILGIWAAGAAYVPLDPNFP.[1].ERLQNIIED 576  Acinetobacter baum...
CAA11794     2563 DQSLSYAELDARANQLARWLIEQG.[4].DLVAVM.[3].SLELVIALLAATKTGGAWLPIDPDYP.[1].DRIAYMLDD 2635 Amycolatopsis orie...
O68007       1573 DQMLTYRELNEKANQLARTLRQKG.[4].SVVGIM.[3].SLEMLTGILAVLKAGGAYMPIDPGLP.[1].ERIQYLITD 1645 Bacillus lichenifo...
Q70LM7       1243 EQRVTYRELNERVNQLAHTLREKG.[4].DLVMLM.[3].SVEMMVAIFAVLKAGGAYLPIDPHSP.[1].ERIAYIFAD 1315 Brevibacillus para...
WP_012265835   29 TCPLTYQQLDHLSNQVAAYLQTQG.[4].TRVGIM.[3].NPGMIIGILGILKAGGCYVPLDPDYP.[1].ERLRYILDH 101  Microcystis aerugi...
NP_252017     469 GGTLSYAELDAKVQAVADALRAAG.[4].ERVALL.[3].GPHLLPAILGVQRAGGAYVPINPDHP.[1].ERVRLLLED 541  Pseudomonas aerugi...
O68008       3023 KEELSYKALNERSNQLAGLLREKG.[4].MIVGVM.[3].SVEMIVGMLAVLKAGGAYLPIDPEYP.[1].DRIRYMIED 3095 Bacillus lichenifo...

Feature 2                                        #   ###### ##
query          89 .[2].ATLVLA.[40].R       REDPAYIIYTSGSTGTPKGVVVSHGALAGHCRAVVD.[7].DAVLQFAS.[3].DT 193
1AMU_A        135 .[2].ARMLLT       Q.[37].STDLAYVIYTSGTTGNPKGTMLEHKGISNLKVFFEN.[7].DRIGQFAS.[3].DA 236  Brevibacillus brevis
3VNQ_A        131 .[2].ASALIG       E.[34].AEDMAYVIYTSGTTGNPKGVPVRHANVLALLAGAPS.[7].DRWLLFHS.[3].DF 229  Streptomyces sp.
4ZXH_A        577 .[2].PKVILT       Q.[32].RGDIAYVMYTSGSTGKPKGVRIGHPSIINFLLSMND.[7].TQLLAITT.[3].DI 673  Acinetobacter baum...
CAA11794     2636 .[2].PALVIT       T.[48].RRHPAYLIYTSGSTGRPKAVVITHRNLTNYLFHCGR.[6].GRSVMHSS.[3].DL 2747 Amycolatopsis orie...
O68007       1646 .[2].ADLLLT       Q.[34].RGDLAYVIYTSGTTGNPKGVMVEHRNIIHAHYTWRK.[8].VNLLQLAS.[3].DV 1745 Bacillus lichenifo...
Q70LM7       1316 .[2].AKLVLA       Q.[34].RGDLVYVMYTSGSTGKPKGVMIEHGALLNVLHGMQD.[7].DAFLLKTT.[3].DI 1414 Brevibacillus para...
WP_012265835  102 .[2].IEILLT       E.[56].REDLMVILYTSGSTGRPKGVMLNHRGYMNRLEWMQK.[7].DRVAQRTS.[3].DI 222  Microcystis aerugi...
NP_252017     542 .[2].ARVVLV       D.[34].RGDLAYVIYTSGSTGMPKGVMVEHRSVVNRLNWMQR.[7].DVLLQKTP.[3].DV 640  Pseudomonas aerugi...
O68008       3096 .[2].ISILLK       K.[33].SSDMAYVIYTSGSTGKPKGVMVNHQSIVNTLYWRKQ.[7].DATLQVPS.[3].DS 3193 Bacillus lichenifo...

Feature 2
query         194 ALEQILPALA.[1].GARLVIGG.[8].FLKLLER.[2].ITVADLPPVYLRE.[15].SLRLLICGGE.[2].GPDVVA 275
1AMU_A        237 SVWEMFMALL.[1].GASLYIIL.[9].FEQYINQ.[2].ITVITLPPTYVVH.[ 7].SIQTLITAGS.[2].SPSLVN 311  Brevibacillus brevis
3VNQ_A        230 SVWEIWGAFS.[1].GAELVVLP.[9].YLAVIID.[2].VTVINQTPTAFLA.[13].GLRYVIFGGE.[2].TAPMLR 310  Streptomyces sp.
4ZXH_A        674 SILELLIPLM.[1].GGVVHVCP.[9].LVDYLNA.[2].INVLQATPATWKM.[ 9].AGLTALCGGE.[2].DTILAE 750  Acinetobacter baum...
CAA11794     2748 TITAMFTPLT.[1].GGTVHVGA.[3].VIGAVDS.[2].SIFLKATPSHLRT.[ 9].VSGDLLLGGE.[2].PVDTIV 2818 Amycolatopsis orie...
O68007       1746 FAGDLCRSLL.[1].GGTMYIVP.[9].LYDMINK.[2].IHMLESTPSLIIP.[13].SMKLLIMGSD.[6].YKWLVE 1830 Bacillus lichenifo...
Q70LM7       1415 SVAEIFGWVP.[2].GKLVILEP.[8].IWQAVVG.[2].ITHINFVPSMLIP.[12].RLRYILACGE.[2].PDELVP 1494 Brevibacillus para...
WP_012265835  223 SVWEIFWTLM.[1].GATICPVK.[9].FAAWIKK.[2].INVMHFVPSLFGE.[13].DLRWLIFSGE.[2].PMSFIQ 303  Microcystis aerugi...
NP_252017     641 SVWELFWWSF.[1].GARLSLLP.[9].MLRSIQR.[2].VTVIHFVPSMLTP.[16].SLRLVFCSGE.[2].APLQVA 724  Pseudomonas aerugi...
O68008       3194 SVEDIFTTLI.[1].GAKLVLIR.[8].IIGVLRT.[2].ATNLLAVPSFYLN.[10].DLRFVTVAGE.[2].NESLIR 3270 Bacillus lichenifo...

Feature 2
query         276 LW.[8].RLLNAYGPTEAT.[2].ALVHDV.[ 8].VPIGRPL.[2].TRIYILD.[11].GELHIGG.[2].LAIGYHG 356
1AMU_A        312 KW.[4].TYINAYGPTETT.[2].ATTWVA.[ 8].VPIGAPI.[2].TQIYIVD.[11].GELCIGG.[2].LARGYWK 388  Brevibacillus brevis
3VNQ_A        311 PW.[9].RLVNGYGITETT.[2].TTFEEI.[ 9].SIIGRAL.[2].FGTRVVG.[11].GELWLSG.[2].LAEGYLR 393  Streptomyces sp.
4ZXH_A        751 KL.[5].CLWNVYGPTETT.[2].SSAARI.[ 5].IDLGEPL.[2].TQLYVLD.[11].GELWIGG.[2].LAVDYWQ 825  Acinetobacter baum...
CAA11794     2819 QW.[7].VVVNEYGPTEAT.[2].CVEYRL.[11].VPIGTPL.[2].MRAFVLD.[11].GELYVSG.[2].VARGYLG 2901 Amycolatopsis orie...
O68007       1831 RF.[4].RIINSYGVTEAS.[2].SGYYEE.[10].TPIGKPL.[2].TAFYILD.[11].GELYIGG.[2].IARGYLN 1909 Bacillus lichenifo...
Q70LM7       1495 KV.[7].KLENIYGPTEAT.[2].ASRYSL.[ 8].VPIGKPL.[2].YRMYIIN.[11].GELCIAG.[2].LARGYLN 1574 Brevibacillus para...
WP_012265835  304 KW.[8].GLANLYGPTEAS.[2].VTYHII.[11].IPIGKAI.[2].VYLKVLD.[11].GELWLGG.[2].LALGYLK 387  Microcystis aerugi...
NP_252017     725 RF.[8].RLVNLYGPTEAT.[2].VSDHEC.[ 7].VPIGRPI.[2].LRLYVLD.[11].GELYIGG.[2].VARGYLN 804  Pseudomonas aerugi...
O68008       3271 QH.[7].KLFNEYGPTENS.[2].STRGEL.[ 6].VVIGRPI.[2].HKVYILN.[11].GELCLSG.[2].LARGYLN 3348 Bacillus lichenifo...

Feature 2
query         357 RDDLTRERF.[12].RLYRTGDLAAFL.[4].GTVAFHGRIDDQVKIR 409
1AMU_A        389 RPELTSQKF.[ 9].KLYKTGDQARWL.[2].GNIEYLGRIDNQVKIR 436  Brevibacillus brevis
3VNQ_A        394 RPELTAEKF.[12].RYYRTGDLVSEL.[2].GRFAYEGRADLQIKLR 444  Streptomyces sp.
4ZXH_A        826 RPELTDAQF.[10].RLYRTGDKVCLR.[2].GRLTHHGRLDFQVKIR 874  Acinetobacter baumannii AB307-0294
CAA11794     2902 RAGLTASRF.[ 9].RMYRTGDLVRWN.[2].GQLVFAGRVDDQVKVR 2949 Amycolatopsis orientalis
O68007       1910 KPELTKERF.[ 9].NMYKTGDLARWL.[2].GNVEFLGRIDHQVKIR 1957 Bacillus licheniformis
Q70LM7       1575 NPALTEEKF.[ 9].RIYRTGDLARYR.[2].GNIEYLGRMDHQVKIR 1622 Brevibacillus parabrevis
WP_012265835  388 DPEKTAKAF.[11].YIYRTGDLVKEL.[2].GTLEYHGRIDNMVKIR 437  Microcystis aeruginosa
NP_252017     805 RPELNAERF.[ 9].RLYRTGDLARWL.[2].GNLEYLGRADDQVKIR 852  Pseudomonas aeruginosa
O68008       3349 RPDLTLEKF.[ 9].SMYRTGDLARFL.[2].GQIEYLGRIDHQVKIR 3396 Bacillus licheniformis
```

Fig. S8: Alignment of the amino acid sequence of the A domain from the last module of sbnQ from S. indica PHM037 with the reference domains of the CDD search tool. The active site residues are highlighted in yellow. The residues just before the active site (red box) do not match the reference sequences in length.

```
Feature 2
query           7 PDAVAVT.[2].AETLTYRRLSRRANRLARHLRQHG.[4].DKVAVL.[3].GGDSIIALLACLTLGAIWVPLDTEAP 78
1AMU_A         53 PNNVAIV.[2].NEQLTYHELNVKANQLARIFIEKG.[4].TLVGIM.[3].SIDLFIGILAVLKAGGAYVPIDIEYP 124  Brevibacillus brevis
3VNQ_A         49 PERTALS.[2].DDRISYGRLDAWSDAVARTLLAEG.[4].DRVALR.[3].GAEAIVAILAILKCGAAYVPVDLRNP 120  Streptomyces sp.
4ZXH_A        495 GDNHALT.[2].DHTVSYRELGQHIAGIAEYLRAHG.[4].DRVGLM.[3].TALLPAAILGIWAAGAAYVPLDPNFP 566  Acinetobacter bauman...
CAA11794     2554 PEATAAI.[2].DQSLSYAELDARANQLARWLIEQG.[4].DLVAVM.[3].SLELVIALLAATKTGGAWLPIDPDYP 2625 Amycolatopsis orient...
O68007       1564 PNHPAAV.[2].DQMLTYRELNEKANQLARTLRQKG.[4].SVVGIM.[3].SLEMLTGILAVLKAGGAYMPIDPGLP 1635 Bacillus licheniformis
Q70LM7       1234 PDKTALV.[2].EQRVTYRELNERVNQLAHTLREKG.[4].DLVMLM.[3].SVEMMVAIFAVLKAGGAYLPIDPHSP 1305 Brevibacillus parabr...
P39845        472 PERLAIR.[2].GGSLTYAELDMYASRLAAHLAARG.[4].SIVGVL.[3].SPDMLIAVLAVLKAGGAYLPLDPAYP 543  Bacillus subtilis su...
WP_012265835   20 PDAIAVL.[2].TCPLTYQQLDHLSNQVAAYLQTQG.[4].TRVGIM.[3].NPGMIIGILGILKAGGCYVPLDPDYP 91   Microcystis aeruginosa
NP_252017     459 PQRTALL.[3].GGTLSYAELDAKVQAVADALRAAG.[4].ERVALL.[3].GPHLLPAILGVQRAGGAYVPINPDHP 531  Pseudomonas aeruginosa

Feature 2                                               #  ###### ##
query          79 .[1].RRLAAMMAD.[2].PRAVVVD.[34].PADAAYMIYTSGTTGTPKGVVVTQRALADHVQAIGA.[7].DVVLHFA 181
1AMU_A        125 .[1].ERIQYILDD.[2].ARMLLTQ.[37].STDLAYVIYTSGTTGNPKGTMLEHKGISNLKVFFEN.[7].DRIGQFA 230  Brevibacillus brevis
3VNQ_A        121 .[1].SRSDFILAD.[2].ASALIGE.[34].AEDMAYVIYTSGTTGNPKGVPVRHANVLALLAGAPS.[7].DRWLLFH 223  Streptomyces sp.
4ZXH_A        567 .[1].ERLQNIIED.[2].PKVILTQ.[32].FGDIAYVMYTSGSTGKPKGVRIGHPSIINFLLSMND.[7].TQLLAIT 667  Acinetobacter baum...
CAA11794     2626 .[1].DRIAYMLDD.[2].PALVITT.[48].PRHPAYLIYTSGSTGRPKAVVITHRNLTNYLFHCGR.[6].GRSVMHS 2741 Amycolatopsis orie...
O68007       1636 .[1].ERIQYLITD.[2].ADLLLTQ.[34].PGDLAYVIYTSGTTGNPKGVMVEHRNIIHAHYTWRK.[8].VNLLQLA 1739 Bacillus lichenifo...
Q70LM7       1306 .[1].ERIAYIFAD.[2].AKLVLAQ.[34].PGDLVYVMYTSGSTGKPKGVMIEHGALLNVLHGMQD.[7].DAFLLKT 1408 Brevibacillus para...
P39845        544 .[1].ERLSYMLKD.[2].ASLLLTQ.[34].GGSLAYVIYTSGSTGQPKGVAVEHRQAVSFLTGMQH.[7].DIVMVKT 646  Bacillus subtilis ...
WP_012265835   92 .[1].ERLRYILDH.[2].IEILLTE.[56].PEDLMVILYTSGSTGRPKGVMLNHRGYMNRLEWMQK.[7].DRVAQRT 216  Microcystis aerugi...
NP_252017     532 .[1].ERVRLLLED.[2].ARVVLVD.[34].PGDLAYVIYTSGSTGMPKGVMVEHRSVVNRLNWMQR.[7].DVLLQKT 634  Pseudomonas aerugi...

Feature 2
query         182 P.[3].DTALEQILPTLV.[1].GARL     LVR.[9].LRRVLTE.[2].VSVADLPPAYLRE.[11].SLRLLIVGG 256
1AMU_A        231 S.[3].DASVWEMFMALL.[1].GASL.[1].IIL.[9].FEQYINQ.[2].ITVITLPPTYVVH.[ 7].SIQTLITAG 302  Brevibacillus brevis
3VNQ_A        224 S.[3].DFSVWEIWGAFS.[1].GAEL.[1].VLP.[9].YLAVIID.[2].VTVINQTPTAFLA.[13].GLRYVIFGG 301  Streptomyces sp.
4ZXH_A        668 T.[3].DISILELLIPLM.[1].GGVV.[1].VCP.[9].LVDYLNA.[2].INVLQATPATWKM.[ 9].AGLTALCGG 741  Acinetobacter baum...
CAA11794     2742 S.[3].DLTITAMFTPLT.[1].GGTV.[1].VGA.[3].VIGAVDS.[2].SIFLKATPSHLRT.[ 9].VSGDLLLGG 2809 Amycolatopsis orie...
O68007       1740 S.[3].DVFAGDLCRSLL.[1].GGTM.[1].IVP.[9].LYDMINK.[2].IHMLESTPSLIIP.[13].SMKLLIMGS 1817 Bacillus lichenifo...
Q70LM7       1409 T.[3].DISVAEIFGWVP.[2].GKLV.[1].LEP.[8].IWQAVVG.[2].ITHINFVPSMLIP.[12].RLRYILACG 1485 Brevibacillus para...
P39845        647 S.[3].DASVWQLFWWSL.[1].GASA.[1].LLP.[9].IVQAIHQ.[2].VTTAHFIPAMLNS.[14].SLKRVFAGG 725  Bacillus subtilis ...
WP_012265835  217 S.[3].DISVWEIFWTLM.[1].GATI.[1].PVK.[9].FAAWIKK.[2].INVMHFVPSLFGE.[13].DLRWLIFSG 294  Microcystis aerugi...
NP_252017     635 P.[3].DVSVWELFWWSF.[1].GARL.[1].LLP.[9].MLRSIQR.[2].VTVIHFVPSMLTP.[16].SLRLVFCSG 715  Pseudomonas aerugi...

Feature 2
query         257 E.[2].AADTVRLW.[8].RLLNAYGPTEAT.[2].CLLHEV.[ 9].IPIGTPL.[2].TRVAIVD.[11].GELLVGG 338
1AMU_A        303 S.[2].SPSLVNKW.[4].TYINAYGPTETT.[2].ATTWVA.[ 8].VPIGAPI.[2].TQIYIVD.[11].GELCIGG 379  Brevibacillus brevis
3VNQ_A        302 E.[2].TAPMLRPW.[9].RLVNGYGITETT.[2].TTFEEI.[ 9].SIIGRAL.[2].FGTRVVG.[11].GELWLSG 384  Streptomyces sp.
4ZXH_A        742 E.[2].DTILAEKL.[5].CLWNVYGPTETT.[2].SSAARI.[ 5].IDLGEPL.[2].TQLYVLD.[11].GELWIGG 816  Acinetobacter baum...
CAA11794     2810 E.[2].PVDTIVQW.[7].VVVNEYGPTEAT.[2].CVEYRL.[11].VPIGTPL.[2].MRAFVLD.[11].GELYVSG 2892 Amycolatopsis orie...
O68007       1818 D.[6].YKWLVERF.[4].RIINSYGVTEAS.[2].SGYYEE.[10].TPIGKPL.[2].TAFYILD.[11].GELYIGG 1900 Bacillus lichenifo...
Q70LM7       1486 E.[2].PDELVPKV.[7].KLENIYGPTEAT.[2].ASRYSL.[ 8].VPIGKPL.[2].YRMYIIN.[11].GELCIAG 1565 Brevibacillus para...
P39845        726 E.[2].APRTAARF.[7].SLIHGYGPTEAT.[2].AAFYVL.[10].IPIGKPV.[2].ARLYVLD.[11].GELYIAG 807  Bacillus subtilis ...
WP_012265835  295 E.[2].PMSFIQKW.[8].GLANLYGPTEAS.[2].VTYHII.[11].IPIGKAI.[2].VYLKVLD.[11].GELWLGG 378  Microcystis aerugi...
NP_252017     716 E.[2].APLQVARF.[8].RLVNLYGPTEAT.[2].VSDHEC.[ 7].VPIGRPI.[2].LRLYVLD.[11].GELYIGG 795  Pseudomonas aerugi...

Feature 2
query         339 .[2].LAQGYHHRPDLTAERF.[ 7].RMYRTGDLASFI.[4].GSIAFHGRLDHQVKIR 395
1AMU_A        380 .[2].LARGYWKRPELTSQKF.[ 9].KLYKTGDQARWL.[2].GNIEYLGRIDNQVKIR 436  Brevibacillus brevis
3VNQ_A        385 .[2].LAEGYLRRPELTAEKF.[12].RYYRTGDLVSEL.[2].GRFAYEGRADLQIKLR 444  Streptomyces sp.
4ZXH_A        817 .[2].LAVDYWQRPELTDAQF.[10].RLYRTGDKVCLR.[2].GRLTHHGRLDFQVKIR 874  Acinetobacter baumannii AB307-0294
CAA11794     2893 .[2].VARGYLGRAGLTASRF.[ 9].RMYRTGDLVRWN.[2].GQLVFAGRVDDQVKVR 2949 Amycolatopsis orientalis
O68007       1901 .[2].IARGYLNKPELTKERF.[ 9].NMYKTGDLARWL.[2].GNVEFLGRIDHQVKIR 1957 Bacillus licheniformis
Q70LM7       1566 .[2].LARGYLNNPALTEEKF.[ 9].RIYRTGDLARYR.[2].GNIEYLGRMDHQVKIR 1622 Brevibacillus parabrevis
P39845        808 .[2].VARGYLNRPALTEERF.[ 9].RMYKTGDVARWL.[2].GNVEFLGRTDDQVKIR 864  Bacillus subtilis subsp. subtilis str....
WP_012265835  379 .[2].LALGYLKDPEKTAKAF.[11].YIYRTGDLVKEL.[2].GTLEYHGRIDNMVKIR 437  Microcystis aeruginosa
NP_252017     796 .[2].VARGYLNRPELNAERF.[ 9].RLYRTGDLARWL.[2].GNLEYLGRADDQVKIR 852  Pseudomonas aeruginosa
```

Fig. S9: Alignment of the amino acid sequence of the A domain from the last module of sbnQ from M. gryphiswaldense with the reference domains of the CDD search tool. The active site residues are highlighted in yellow.

Table S4: Putative open reading frames (ORFs) encoding polyketide synthases (PKSs), non-ribosomal peptide synthetases (NRPSs), or hybrid in PKS and NRPS gene clusters from different magnetotactic bacteria.

| **Species** | **strain** | **Class** | **PKS** | **NRPS** | **Hybrid** | **Trans-AT PKS** |
|---|---|---|---|---|---|---|
| ***Magnetospirillum gryphiswaldense*** | MSR-1 | α | 0 | 0 | 0 | 1 Trans-AT PKS |
| ***Magnetospirillum magneticum*** | AMB-1 | α | 1 T1PKS | 1 NRPS-like | 0 | 0 |
| ***Magnetospira*** **sp.** | QH-2 | α | 0 | 1 NRPS | 0 | 0 |
| ***Magnetospira*** **sp.** | ME-1 | α | 1 T1PKS | 1 NRPS-like | 0 | 0 |
| ***Magnetovibrio blakemorei*** | MV-1 | α | 0 | 0 | 0 | 1 Trans-AT PKS |
| ***Magnetospirillum*** **sp.** | XM-1 | α | 1 T1PKS | 1 NRPS-like | 0 | 0 |
| ***Magnetofaba australis*** | IT-1 | α | 1 T1PKS | 0 | 0 | 0 |
| ***Magnetospirillum magnetotacticum*** | MS-1 | α | 1 T1PKS | 1 NRPS-like | 0 | 0 |
| ***Magnetospirillum*** **sp.** | SO-1 | α | 1 T1PKS | 1 NRPS-like | 0 | 0 |
| ***Magnetospirillum marisnigri*** | SP-1 | α | 1 T3PKS | 1 NRPS-like | 0 | 0 |
| ***Magnetospirillum*** **sp.** | 15-1 | α | 1 T1PKS | 1 NRPS-like | 0 | 1 Trans-AT PKS |
| **Magnetospirillum sp.** | 64-120 | α | 0 | 0 | | 1 Trans-AT PKS |
| ***Magneto*****-ovoid bacterium** | MO-1 | α | 0 | 1 NRPS / 1 NRPS-like | 0 | 0 |
| ***Desulfovibrio magneticus*** | RS-1 | δ | 0 | 1 NRPS-like | 0 | 0 |
| ***Desulfamplus magnetovallimortis*** | BW-1 | δ | 2 T1PKS | 0 | 0 | 0 |
| ***Candidatus*** **Magnetoglobus multicellularis str.** | Araruama | δ | 1 T1PKS | 5 NRPS / 2 NRPS-like | | 2 Trans-AT PKS like |
| **Ectothiorhodospiraceae bacterium** | BW-2 | γ | T2PKS | 1 NRPS-like | 2 NRPS/T1PKS | 0 |
| **Gamma Proteobacterium** | SS-5 | γ | 0 | 1 NRPS | 2 NRPS/T1PKS | 1 Trans-AT PKS |
| ***Candidatus*** **Magnetobacterium casensis** | MYR-1 | Nitrospirae | 0 | 1 NRPS-like | 0 | 0 |
| ***Candidatus*** **Magnetobacterium bavaricum** | TM-1 | Nitrospirae | 0 | 1 NRPS-like | 0 | 0 |
| ***Candidatus*** **Magnetoovum chiemensis** | CS-04 | Nitrospirae | 1 T1PKS | 1 NRPS | 0 | 0 |

Table S5: Strains and vectors used in this study.

| **Strain or vector** | **Relevant characteristic (s)** | **Reference and/or source** |
|---|---|---|
| Strains | | |
| *E. coli* | | |
| DH5α | Host for cloning; $F^+$ *endA1 glnV44 thi-1 recA1 relA1 gyrA96 deoR nupG purB20* φ80d*lac*ZΔM15 Δ(*lacZYA-argF*)U169, hsdR17($r_K^- m_K^+$), $\lambda^-$ | (1) |
| WM3064 | Conjugation strain; *thrB1004 pro thi rpsL hsdS lac*ZΔM15 *RP4-1360* Δ(*araBAD*)*567* Δ*dapA1341*::[*erm pir*] | William Metcalf at UIUC |
| *M. gryphiswaldense* | | |
| Wild type | MSR-1 R3/S1; $Rif^R$, $Sm^R$ | (2) |
| Δ*trans-at-pks* | Core-biosynthetic genes of trans-AT PKS deletion strain (MSR-1_15620-15650) | This study |
| $P_{mamDC45}$-*trans-at-pks* | Strain with chromosomally inserted $P_{mamDC45}$ promoter in front of MSR-1_15600 | This study |
| Vectors | | |
| pORFM | universal in-frame deletion/in-frame fusion vector for GalK-based counterselection; *npt galK tetR mob*RK2 | (3) |
| pORFM-Δ*trans-at-pks* | Vector for chromosomal deletion of core-biosynthetic genes of trans-AT PKS gene cluster (MSR-1_15620-15650) | This study |
| pORFM-$P_{mamDC45}$-*trans-at-pks* | Vector for insertion of a promoter 1xPmamDC45-oRBS in front of MSR-1_15600 | This study |

Table S6: Primers used in this study.

| No. (RPA) | Primer name | Sequence 5’-3’ |
|---|---|---|
| ***Site-specific chromosomal deletion/insertion by homologous recombination*** | | |
| 595 | F1_del_Trnpks-nrps | gtcattactggatctatcaacaggagtcctgcagtaggatgagcatcgccgctttcctggg |
| 596 | R1_del_Trnpks-nrps | gctggatcggttagcccgaggctttcatgcatggcctccttcgc |
| 597 | F2_del_Trnpks-nrps | ggaggccatgcatgaaagcctcgggctaaccgatccagcataatatg |
| 598 | R2_del_Trnpks-nrps | gcggcagcgtgaagctagcatcactagtctagcgcagcaggtcatcgatggagcgg |
| 599 | sq_Rev1_Trnpks_nrps | cccatacaggcggtcaaacag |
| 600 | sq_For2_Trnpks_nrps | cgaggtggtgttcgtggtc |
| 601 | sq_Rev2_Trnpks_nrps | ggcttttggcgatgatctgc |
| 602 | sq_For3_Trnpks_nrps | accgctacatcatcgtcgac |
| 937 | F1_Pro_in_PKS_NRPS | caggaaagacttaagctgcagtagggcccgggtgatggtcgcc |
| 938 | R1_Pro_in_PKS_NRPS | gagaactaagagctagtaaagcgaaaaagtcttacttgtcttgtccggcg |
| 939 | F2_Pro_in_PKS_NRPS | cgccatcgccggacaagacaagtaagactttttcgctttactagctc |
| 940 | R2_Pro_in_PKS_NRPS | ggtgctgacgagacgaagaacatgcatatgctgatctcctaagcttcgc |
| 941 | F3_Pro_in_PKS_NRPS | ccctgcgaagcttaggagatcagcatatgcatgttcttcgtctcgtcagc |
| 942 | R3_Pro_in_PKS_NRPS | ctctagactaaagcttatcgaattcctagccagaaccgtatagaacaattcg |
| 943 | Ck_barA_CT_R | catgtcgttgccgtcaagc |
| 944 | Ck_P_PKS_NRPS_F | gaccttgtacgaatgctgcc |
| 945 | Ck_P_PKS_NRPS_R | cgcatcaattcgtggtccag |
| 946 | Ck_yojil1_NT_F | gcaccttggaaaatctcggc |
| 484 | sq_bk_pORFM_F | gccactcatcgcagtctagc |
| 485 | sq_pORFM_bk_Rev | tctgcggactggctttctac |

**NMR data and spectra**

Table S7: NMR spectroscopic data for Sesbanimide R in methanol-$d_4$ at 500/125 MHz.

| # | δ $^{13}$C [PPM] | δ $^{1}$H [PPM], MULT (*J* [HZ]) | COSY | HMBC |
|---|---|---|---|---|
| **1** | 174.6 | - | - | - |
| **1a** | 174.6 | - | - | - |
| **2** | 39.1 | 2.36, 2.68, m | 2a, 3 | 1, 3, 4 |
| **2a** | 37.7 | 2.33, 2.70, m | 2, 3 | 1a, 3, 4 |
| **3** | 28.4 | 2.34, m | 2, 2a, 4 | 1, 1a, 2, 2a, 4, 5 |
| **4** | 38.9 | 1.49, 1.73, m | 3, 5 | 2, 2a, 3, 5, 6 |
| **5** | 71.7 | 3.98, dt (10.4, 3.1) | 4, 6 | 3, 4, 6, 7 |
| **6** | 90.6 | 3.66, d (3.1) | 5 | 4, 5, 7, 11 |
| **7** | 213.2 | - | - | - |
| **8** | 46.9 | 3.71, q (7.0) | 12 | 7, 9, 10, 12, 13 |
| **9** | 144.5 | - | - | - |
| **10** | 66.8 | 4.66, s | 13 | 8, 9, 13, 14 |
| **11** | 60.6 | 3.40, s | - | 6 |
| **12** | 16.6 | 1.19, d (7.0) | 8 | 7, 8, 9, 13 |
| **13** | 113.9 | 4.96, 5.18, s | 10 | 8, 9, 10 |
| **14** | 174.6 | - | - | - |
| **15** | 34.8 | 2.38, t (7.5) | 16 | 14, 16, 17 |
| **16** | 26.0 | 1.63, m | 15, 17 | 14, 15, 17, 18 |
| **17** | 30.4 | 1.34, m | 16, 18 | 15, 16, 18, 19 |
| **18** | 30.1 | 1.34, m | 17, 19 | 16, 17, 19, 20 |
| **19** | 30.6 | 1.30, m | 18, 20 | 17, 18, 20, 21 |
| **20** | 29.7 | 1.45, m | 19, 21 | 18, 19, 21, 22 |
| **21** | 33.8 | 2.18, q (7.1) | 20, 22 | 19, 20, 22, 23 |
| **22** | 144.1 | 6.10, dt (7.2, 15.1) | 21, 23 | 20, 21, 23, 24, 25 |
| **23** | 129.7 | 6.22, dd (10.8, 15.1) | 22, 24 | 21, 22, 24, 25 |
| **24** | 142.3 | 7.12, dd (10.7, 15.1) | 23, 25 | 22, 23, 25, 26 |
| **25** | 122.9 | 6.02, d (15.1) | 24 | 22, 23, 24, 26 |
| **26** | 168.3 | - | - | - |
| **27** | 177.6 | - | - | - |
| **28** | 54.9 | 4.40, dd (5.3, 7.7) | nd | 26, 27, 29, 30 |
| **29** | 30.9 | 1.92, m | 30 | 27, 28, 30, 31 |
| **30** | 26.0 | 1.63, m | 29, 31 | 28, 29, 31 |
| **31** | 41.9 | 3.22, m | 30 | 29, 30, 32 |
| **32** | 158.4 | - | - | - |

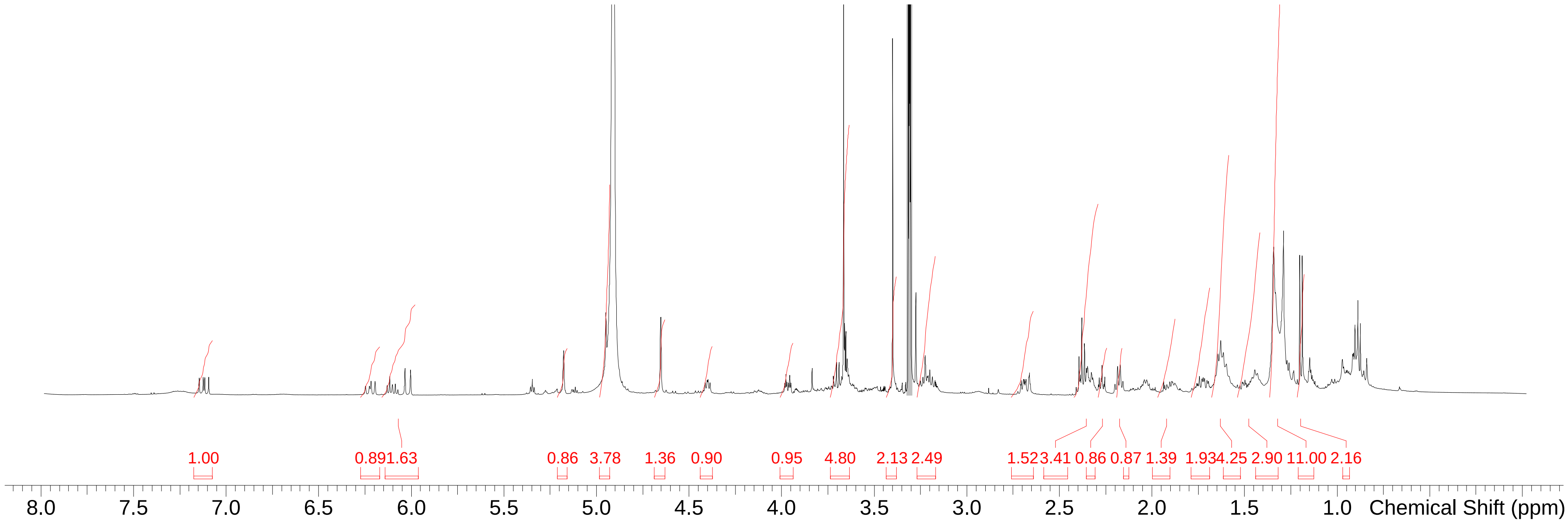


Fig. S10: $^{1}$H spectrum of Sesbanimide R in methanol-$d_4$ at 500 MHz.

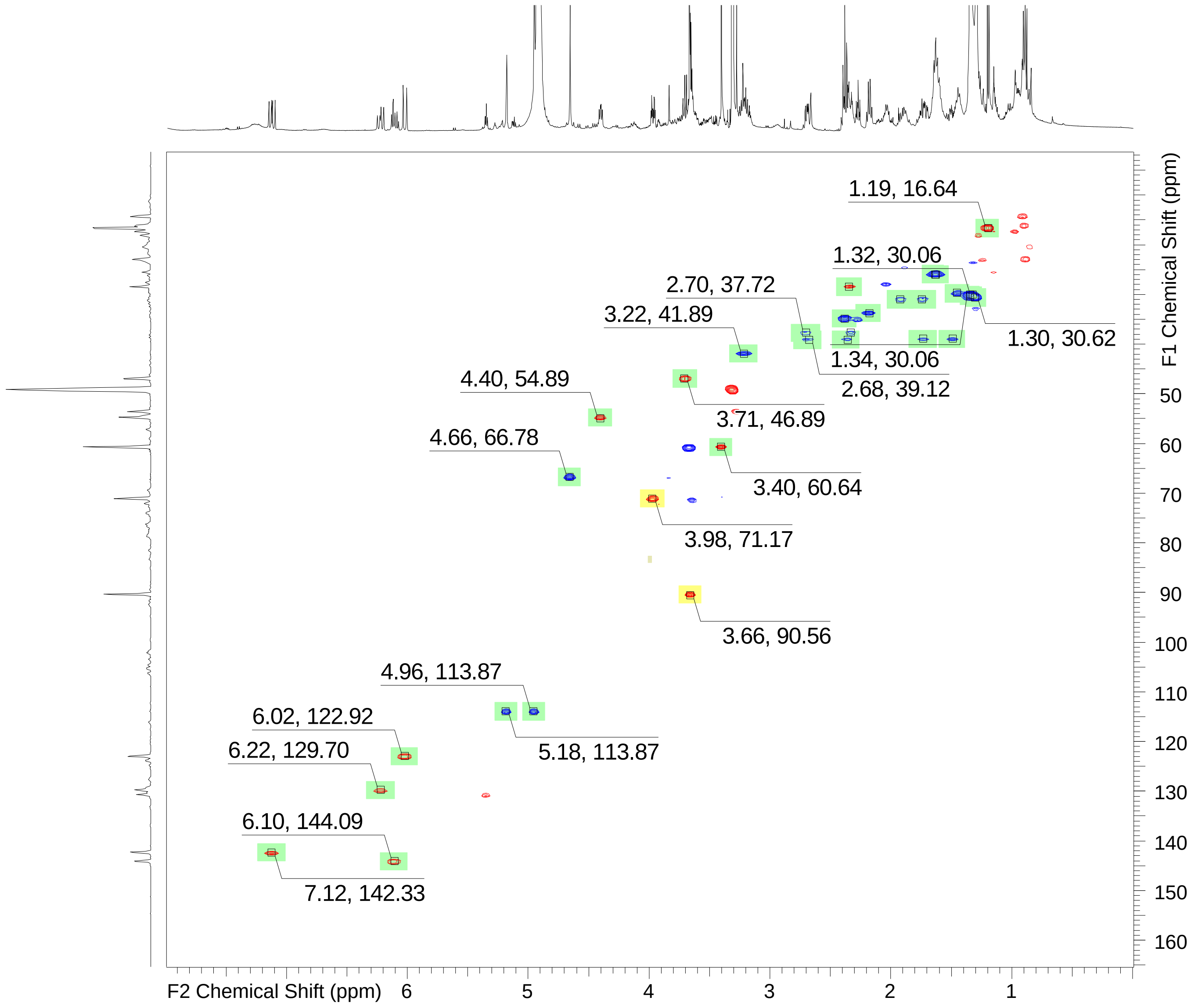


Fig. S11: HSQC spectrum of Sesbanimide R in methanol-$d_4$ at 500/125 MHz.

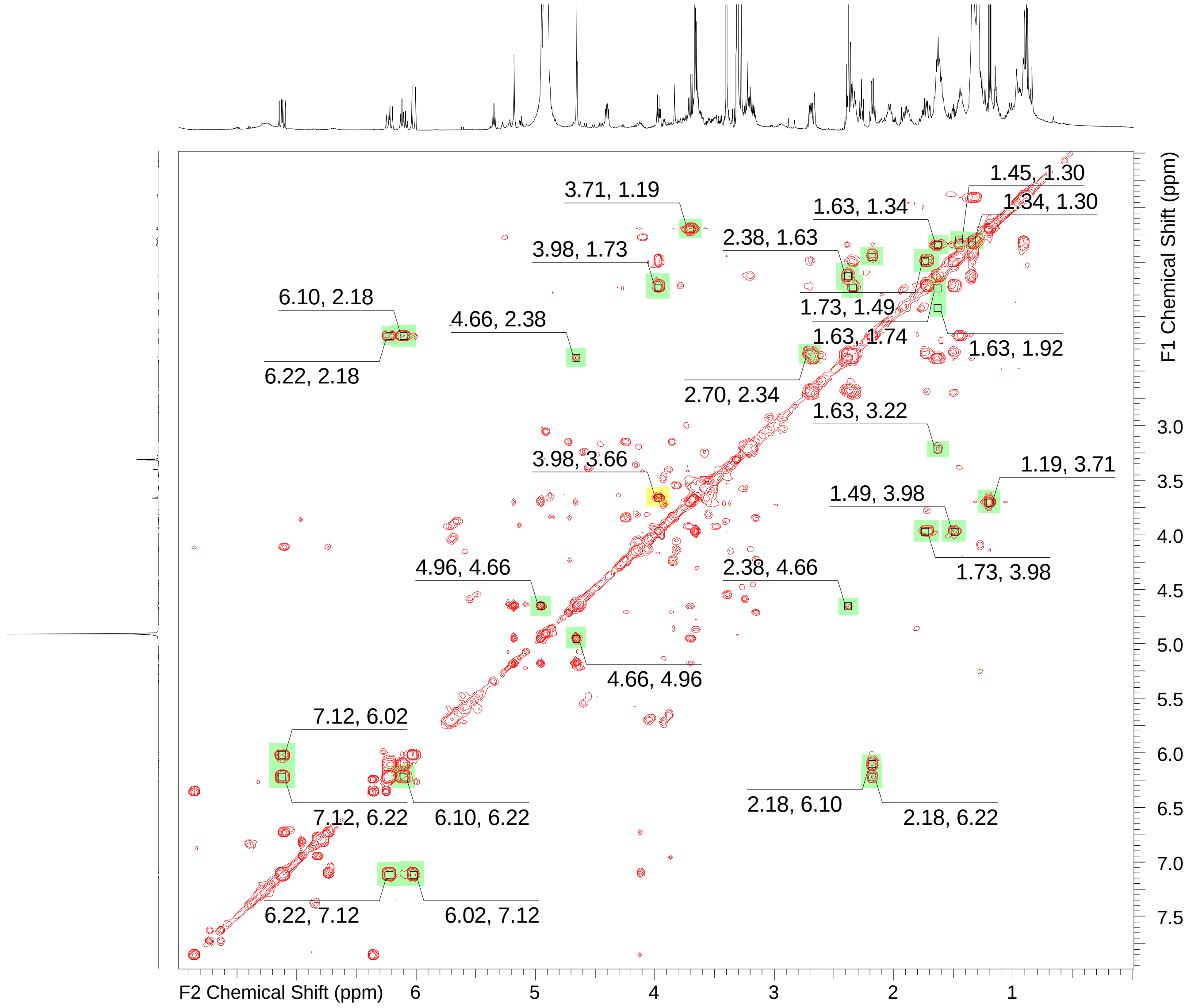


Fig. S12: COSY spectrum of Sesbanimide R in methanol-$d_4$ at 500 MHz.

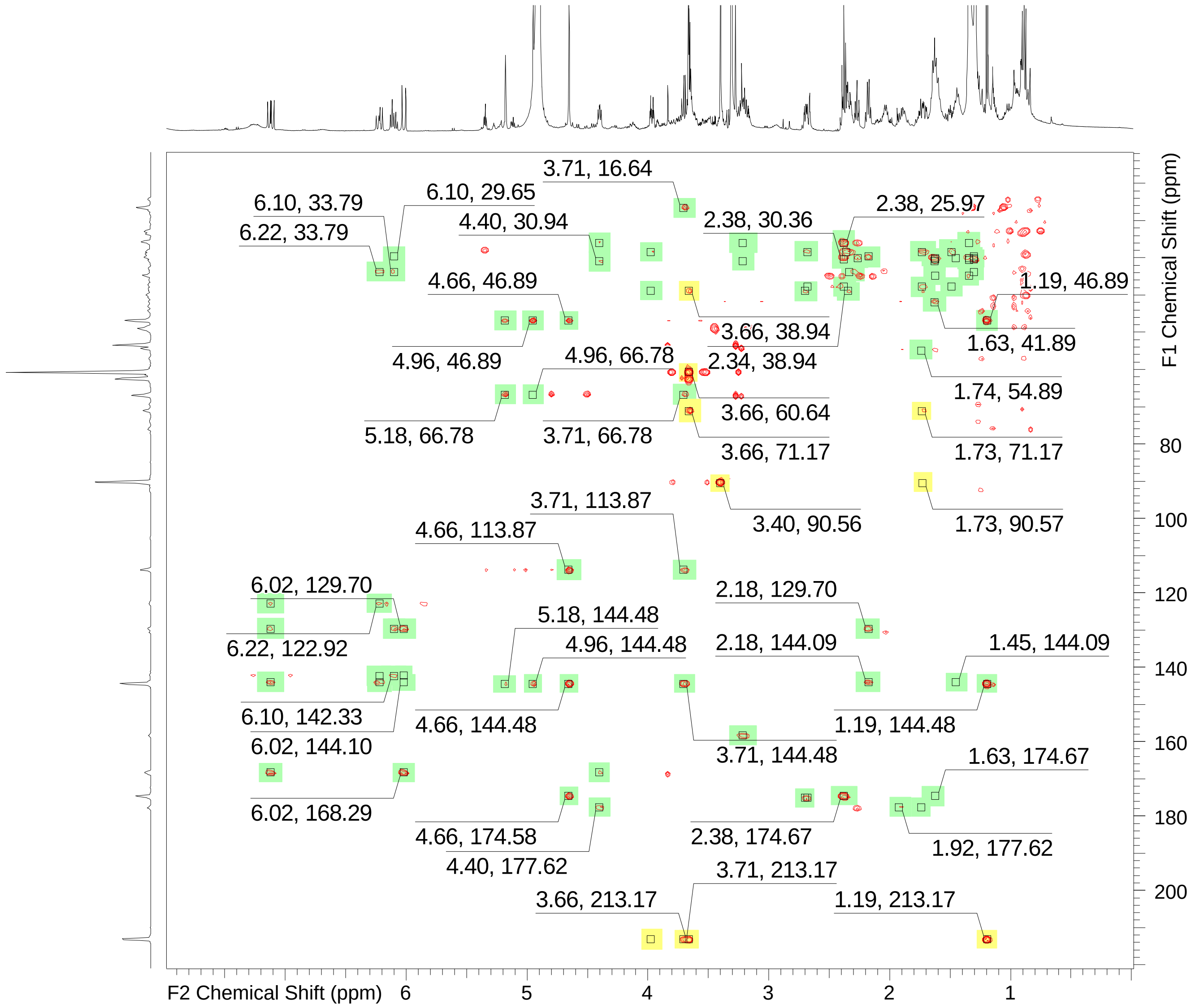


Fig. S13: HMBC spectrum of Sesbanimide R in methanol-$d_4$ at 500/125 MHz.

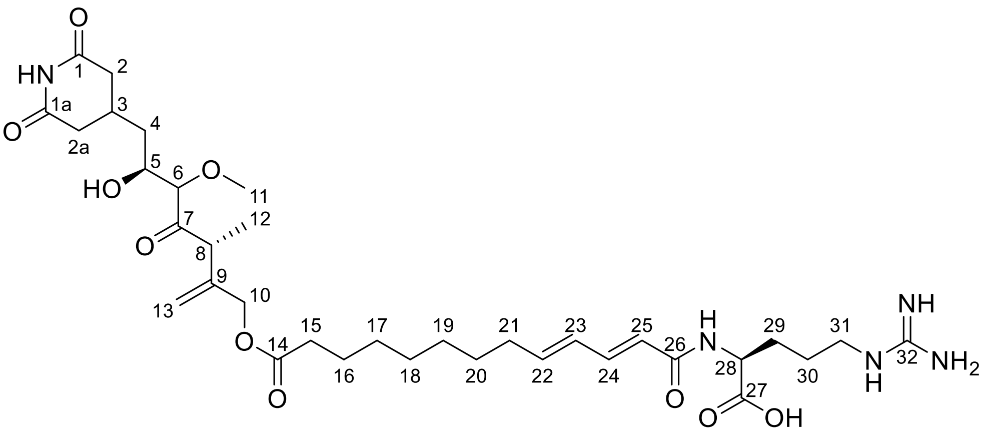


Fig. S14: NMR elucidated structure of sesbanimide R with the carbon atoms numbered from 1 to 32.

References


1. Hanahan D. 1983. Studies on transformation of Escherichia coli with plasmids. Journal of Molecular Biology 166:557–580. doi:10.1016/s0022-2836(83)80284-8.
2. Schultheiss D, Kube M, Schüler D. 2004. Inactivation of the flagellin gene flaA in Magnetospirillum gryphiswaldense results in nonmagnetotactic mutants lacking flagellar filaments. Appl Environ Microbiol 70:3624–3631. doi:10.1128/AEM.70.6.3624-3631.2004.
3. Raschdorf O, Plitzko JM, Schüler D, Müller FD. 2014. A tailored galK counterselection system for efficient markerless gene deletion and chromosomal tagging in Magnetospirillum gryphiswaldense. Appl Environ Microbiol 80:4323–4330. doi:10.1128/AEM.00588-14.
